# Distinct SoxB1 networks are required for naïve and primed pluripotency

**DOI:** 10.1101/229716

**Authors:** Andrea Corsinotti, Frederick C. K. Wong, Tülin Tatar, Iwona Szczerbinska, Florian Halbritter, Douglas Colby, Sabine Gogolok, Raphaël Pantier, Kirsten Liggat, Elham S. Mirfazeli, Elisa Hall-Ponsele, Nicholas Mullin, Valerie Wilson, Ian Chambers

## Abstract

Deletion of *Sox2* from embryonic stem cells (ESCs) causes trophectodermal differentiation. While this can be prevented by enforced expression of the related SOXB1 proteins, SOX1 or SOX3, the roles of SOXB1 proteins in epiblast stem cell (EpiSC) pluripotency are unknown. Here we show that *Sox2* can be deleted from EpiSCs with impunity. This is due to a shift in the balance of SoxB1 expression in EpiSCs, which have decreased Sox2 and increased Sox3 compared to ESCs. Consistent with functional redundancy, *Sox3* can also be deleted from EpiSCs without eliminating self-renewal. However, deletion of both *Sox2* and *Sox3* prevents self-renewal. The overall SOXB1 levels in ESCs affect differentiation choices: neural differentiation of *Sox2* heterozygous ESCs is compromised, while increased SOXB1 levels divert the ESC to EpiSC transition towards neural differentiation. Therefore, optimal SOXB1 levels are critical for each pluripotent state and for cell fate decisions during exit from naïve pluripotency.

## Introduction

Pluripotent cells have the unique ability to differentiate into every cell type of an adult organism (Nichols & Smith, 2009). During mouse development, the embryo contains pluripotent cells in the epiblast until the onset of somitogenesis (Osorno *et al,* 2012; Chambers & Tomlinson, 2009). Distinct pluripotent cell types can be isolated in culture from the preimplantation and postimplantation epiblast. Embryonic stem cells (ESCs) (Evans & Kaufman, 1981; Martin, 1981) from preimplantation embryos are termed ‘naïve’ pluripotent cells. Epiblast stem cells (EpiSCs) (Tesar *et al,* 2007; Brons *et al,* 2007), commonly isolated from the post-implantation epiblast, are known as ‘primed’ pluripotent cells. Naïve and primed cells differ dramatically in responses to extracellular signals (Nichols & Smith, 2009). ESCs self-renew in response to a combination of leukaemia inhibitory factor (LIF) and either foetal calf serum (FCS), bone morphogenic protein (BMP) or Wnt (Smith *et al,* 1988; Ying *et al,* 2003a; ten Berge *et al,* 2011) and differentiate in response to fibroblast growth factor (FGF) (Kunath *et al,* 2007; Stavridis *et al,* 2007). In contrast, FGF together with Nodal/Activin A are required for EpiSCs self-renewal (Vallier *et al,* 2009; Guo *et al,* 2009).

Pluripotency is regulated by a pluripotency gene regulatory network (PGRN) (Chambers & Tomlinson, 2009; Festuccia *et al,* 2013; Wong *et al,* 2016). While some transcription factors such as Esrrb and T (Brachyury) are associated more closely with naïve or primed pluripotent states respectively (Festuccia *et al,* 2012; Osorno *et al,* 2012; Tsakiridis *et al,* 2014), the core TFs of the PGRN, Nanog, Sox2 and Oct4 (the *Pou5f1* gene product, also referred as Oct3/4) are expressed in both naïve and primed pluripotent cells (Niwa *et al,* 2000; Masui *et al,* 2007; Avilion *et al,* 2003; Chambers *et al,* 2003; Karwacki-Neisius *et al,* 2013; Osorno *et al,* 2012; Festuccia *et al,* 2012; Brons *et al,* 2007; Tesar *et al,* 2007). While the role of Sox2 has been extensively characterised in naïve cells (Wong *et al,* 2016), its role in primed pluripotency is less well known.

Sox2 is a member of a family of twenty Sox TFs (Pevny & Lovell-Badge, 1997; Kamachi & Kondoh, 2013). All SOX proteins contain a High-Mobility-Group (HMG) box DNA binding domain closely related to the founding member of the Sox family, SRY (Kondoh & Lovell-Badge, 2016). While some SOX proteins contain a transcriptional activation domain, others contain repression domains (Uchikawa *et al,* 1999; Bowles *et al,* 2000; Ambrosetti *et al,* 2000). The paradigm of action for SOX proteins is that they bind to target gene sequences through a DNA-mediated interaction with a partner protein, to specify target gene selection (Kamachi *et al,* 1999; Remenyi, 2003; Williams *et al,* 2004; Kamachi & Kondoh, 2013). In pluripotent cells the principal interaction of SOX2 with OCT4 (Ambrosetti *et al,* 1997, 2000) is considered to positively regulate expression of many pluripotency-specific genes including *Nanog, Oct4* and *Sox2* (Tomioka *et al,* 2002; Chew *et al,* 2005; Okumura-Nakanishi *et al,* 2005; Rodda, 2005; Kuroda *et al,* 2005). Loss of SOX2 in ESCs induces trophoblast differentiation, phenocopying OCT4 loss and supporting the idea of a mutually dependent mode of action (Niwa *et al,* 2000; Masui *et al,* 2007).

Analysis of sequence conservation within the HMG box has divided the Sox family into eight groups that can be further divided into subgroups based on homology outside the HMG box (Kondoh & Lovell-Badge, 2015; Kamachi, 2016). Together with SOX1 and SOX3, SOX2 belongs to the SOXB1 group, each of which also contain transcriptional activation domains (Uchikawa *et al,* 1999; Ambrosetti *et al,* 2000; Bowles *et al,* 2000; Kondoh & Kamachi, 2010; Ng *et al,* 2012; Kamachi & Kondoh, 2013). SOXB1 proteins bind the same DNA sequence *in vitro* (Kamachi *et al,* 1999; Kamachi, 2016). Previous studies demonstrated that SOXB1 factors are co-expressed during embryonic development and can substitute for each other in different biological systems, both *in vitro* and *in vivo* (Wood & Episkopou, 1999; Niwa *et al,* 2016; Adikusuma *et al,* 2017). Here we investigate the requirements of naïve and primed pluripotent states for SOXB1 expression. Our results indicate that the essential requirement of SOXB1 function for naïve pluripotent cells extends to primed pluripotent cells. SOX3, which is highly expressed in primed pluripotent cells, functions redundantly with SOX2, rendering SOX2 dispensable in these cells. We further provide evidence that critical SOXB1 levels are required to specify the identity of cells exiting the naïve pluripotent state.

## Results

### A fluorescent reporter of SOX2 protein expression

To investigate the expression of Sox2 in pluripotent cells, a live cell reporter that retained Sox2 function was prepared by replacing the *Sox2* stop codon with a T2A-H2B-tdTomato cassette (Figure 1A; Figure 1 – Figure supplement 1A). Correctly targeted cells were identified by Southern analysis and are referred to as E14<u>T</u>g2a-<u>S</u>ox2-td<u>T</u>omato (TST) cells (Figure 1 – Figure supplement 1B). Fluorescence microscopy of targeted cells showed a close correlation between SOX2 and tdTomato levels (Figure 1 – Figure supplement 2). Moreover, tdTomato expression recapitulated the SOX2 expression pattern in chimeric embryos (Figure 1 – Figure supplement 3). Targeted cells also showed the expected morphological differences when cultured in a combination of LIF plus inhibitors of MEK and GSK3β (LIF/2i), in LIF/FCS, in LIF/BMP or after passaging in Activin/FGF (Figure 1A). These results indicate that Sox2-tdTomato reporter (TST) cells behave normally and provide a useful live cell report of Sox2 expression levels.

**Figure 1:**
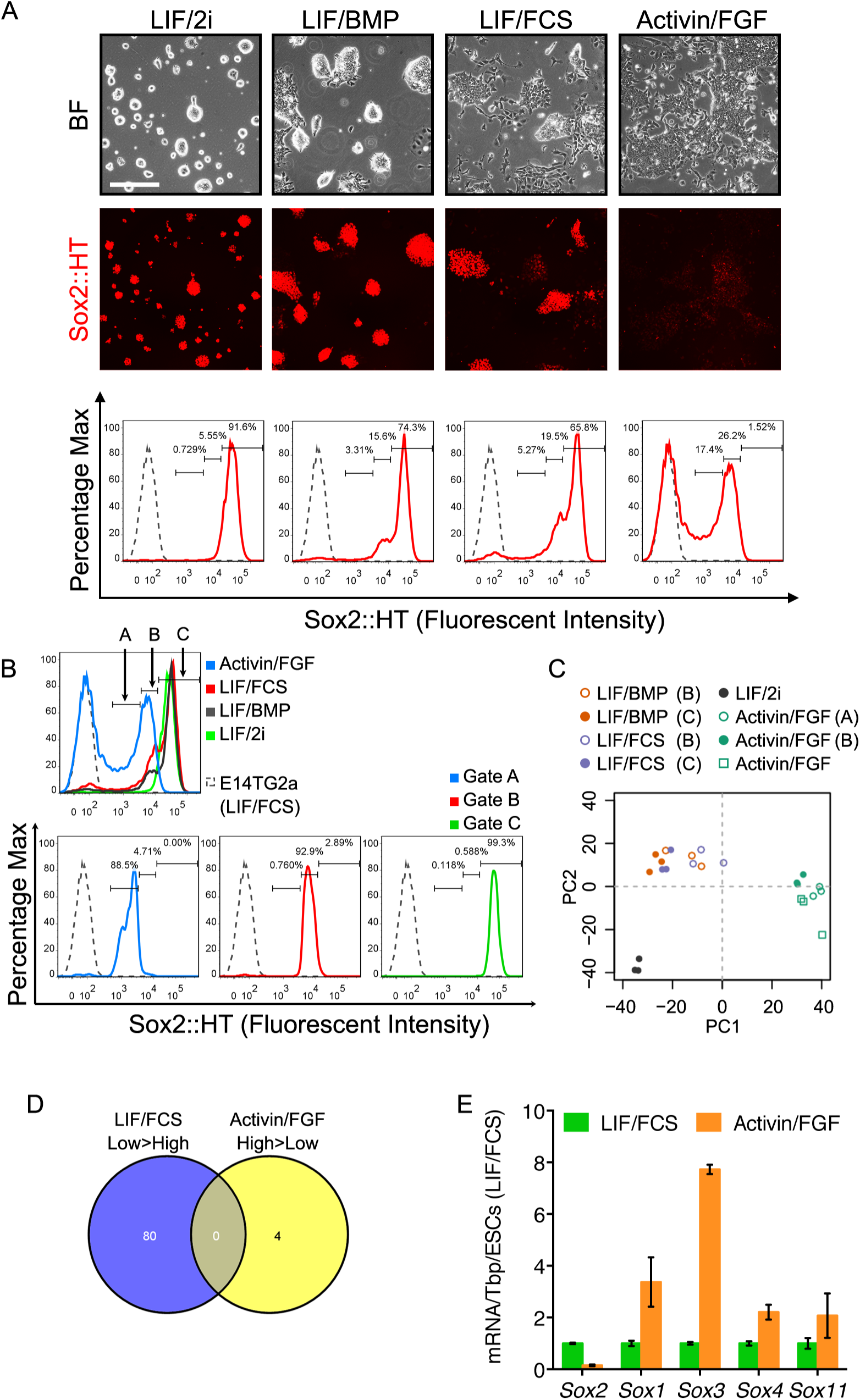
Different roles of Sox2 in preimplantation and postimplantation pluripotency. A. Expression of the Sox2-T2A-H2b-tdTomato (Sox2::HT) reporter from the endogenous *Sox2* allele in targeted TST18 cells. TST18 cells cultured in LIF/FCS/GMEMβ were replated in LIF/2i/N2B27 or LIF/BMP4/N2B27 for 4 passages or in Activin/FGF/N2B27 (Activin/FGF) for 9 passages, examined microscopically (top) and assessed by flow cytometry (bottom); E14TG2a cells were represented as a grey dashed line. B. Three gates (A-C) were used to purify cells for microarray analysis. Gate C captured the Sox2::HT level in LIF/2i cultured ESCs. Gate B captured the overlapping Sox2::HT level in LIF/FCS, LIF/BMP and Activin/FGF cultured cells. Gate A captured the lower Sox2::HT level in Activin/FGF. C. principle component analysis of cells in different culture conditions, either unsorted or sorted using the gates indicated by brackets. D. Despite similar Sox2::HT levels, no differentially expressed genes (DEGs, FDR =0.1) were common to LIF/FCS-low and Activin/FGF-high cell populations. E. RT-qPCR analysis of *Sox1, Sox3, Sox4* and *Sox11* mRNA level in ESCs (LIF/FCS, red bars) and EpiSCs (Activin/FGF, cultured for 16 passages, blue bars). Transcript levels were normalized to TBP and plotted relative to ESCs (horizontal line). Error bars represent standard error of the mean (n=3 to 4).

TST ESCs were next assessed by fluorescence microscopy and FACS. In LIF/2i, tdTomato expression was high and unimodal. In LIF/FCS or LIF/BMP the same predominant high expressing population was present but with a shoulder of reduced expression and a small number of tdTomato-negative cells, which appeared to coincide with morphologically differentiated cells (Figure 1A). In continuous culture in Activin/FGF (p16), tdTomato expression was bimodal with the highest expression levels overlapping the lower expression levels seen in ESCs cultured in LIF/FCS or LIF/BMP (Figure 1A, B).

### Gene expression in ESCs and EpiSCs expressing distinct Sox2 levels

ESCs and EpiSCs expressing distinct Sox2 levels were separated by FACS according to the tdTomato level (Figure 1B). Microarray analysis was then used to compare gene expression in cells from different cultures expressing similar Sox2 levels and in cells from the same cultures expressing distinct Sox2 levels (Supplementary File 1). Cells expressing the highest tdTomato levels in LIF/2i, LIF/FCS or LIF/BMP cultures were purified using gate C. The mid-level expression gate B enabled purification of the Sox2-low fraction of ESCs cultured in LIF/FCS or LIF/BMP as well as the highest Sox2-expressing EpiSCs. EpiSCs were also purified using the lower expression gate A. Re-sorting confirmed effective purification prior to RNA extraction and analysis (Figure 1B). Compared to the Sox2-high population, Sox2-low EpiSCs have upregulated differentiation markers. Transcripts expressed by Sox2-low EpiSCs differed from those expressed by Sox2-low ESCs (Figure 1 – Figure supplement 4; Supplementary File 1) suggesting that ESCs and EpiSCs have distinct differentiation propensities. ESCs from both LIF/BMP and LIF/FCS cultures sorted for highest tdTomato expression (gate C) are enriched for naïve pluripotency transcripts with mRNAs common to both, including Nr5a2, Tbx3 and Tcl1, positively correlated to Sox2 in all samples (Figure 1 – Figure supplement 5). Moreover, principal component analysis indicated that these Sox2-high ESCs cluster together and separately from ESCs cultured in LIF/2i (Figure 1C) as seen by others (Boroviak *et al,* 2014). More importantly, principal component analysis also showed that EpiSCs and ESCs expressing the same Sox2 level (gate B) were transcriptionally distinct (Figure 1C). Moreover, they shared no commonly enriched mRNAs (Figure 1D). Thus Sox2 levels alone do not dictate the distinction between ESC and EpiSC states.

Microarray and RT-qPCR analyses highlighted changes in Sox gene expression levels (Figure 1E, Figure 1 – Figure supplement 6). Although other Sox gene expression changes occurred, the ability of SOXB1 proteins to function redundantly in ESC self-renewal (Niwa *et al* 2016), prompted us to assess the capacity of SOXB1-related proteins to function more widely in pluripotent cells.

### A subset of SOX family proteins can functionally replace *Sox2* in ESC self-renewal

Recently, Niwa *et al.* (2016) reported that ESC self-renewal could be maintained in the absence of SOX2 by expression of other SOXB1 and SOXG proteins (Niwa *et al,* 2016). We addressed the question of functional redundancy with Sox2 in ESCs using an independent *Sox2* conditional knockout (SCKO) ESC line (Favaro *et al,* 2009; Gagliardi *et al,* 2013). SCKO ESCs have one *Sox2* allele replaced by β-geo, the second allele flanked by *loxP* sites and constitutively express a tamoxifen-inducible Cre recombinase (CreER^T2^) (Figure 2a). SCKO ESCs were transfected with a plasmid in which constitutive Sox cDNA expression was linked to hygromycin B resistance and were either left untreated or were simultaneously treated with tamoxifen to excise *Sox2* (Figure 2A). ESC self-renewal was assessed after 8-10 days of hygromycin selection. Transfection of plasmids encoding transcriptional activator proteins of the SOXB1 family (SOX1, SOX2 or SOX3) enabled similar levels of ESC colony formation in the absence of *Sox2* (Figure 2B, C). In contrast, the SOXB2 proteins (SOX14 and SOX21), in which the SOXB1 DNA binding domain is linked to transcriptional repression domains, did not direct undifferentiated ESC colony formation (Figure 2C). Of the SOX proteins tested, only SOX15 showed any capacity to direct undifferentiated ESC colony formation (Figure 2C), as previously described (Niwa *et al,* 2016), possibly because it has the most similar DNA binding domain to SOXB proteins (Kamachi & Kondoh, 2013). These results indicate that ESC self-renewal requires the function of a SOX protein with a SOXB-like DNA binding domain coupled to a transcriptional activating function.

**Figure 2:**
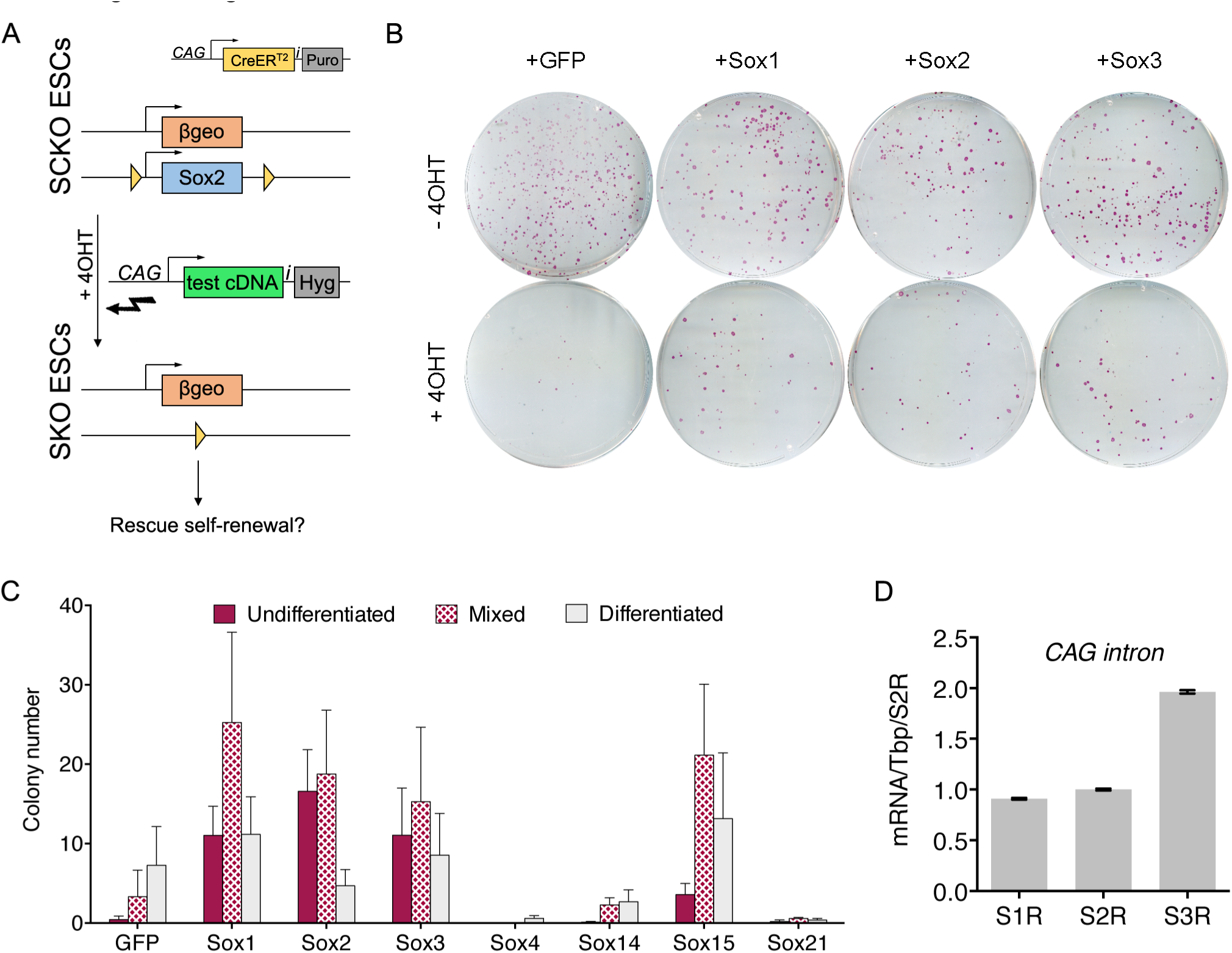
The ESC function of Sox2 can be substituted by SOXB1 and SOXG proteins. A. Strategy for testing the ability of candidate Sox cDNAs to rescue ESC self-renewal upon *Sox2* deletion. *Sox2^fl/-^* (SCKO) ESCs were treated with 4-hydroxy-tamoxifen (4OHT) to induce nuclear localisation of CreER^T2^ and consequent loxP-mediated *Sox2* excision. Simultaneous transfection of test cDNAs linked to hygromycin phosphotransferase via an IRES (i) were tested for Sox2 complementation activity. B. SCKO ESCs transfected with the indicated cDNAs were cultured in the presence of hygromycin B and in the absence (−4OHT) or presence (+4OHT) of 4-hydroxyl-tamoxifen for 8 days before being fixed and stained for alkaline phosphatase (AP) activity. C. Stained colonies were scored based on AP+ (undifferentiated), mixed or AP−(differentiated) morphology. Error bars represent standard error mean (n=3). D. RT-qPCR analysis of the rescuing transgenes of Sox1-, Sox2- and Sox3-rescued (S1R, S2R, S3R) in SCKO ESC populations grown in the presence of 4-hydroxyl-tamoxifen as described in **Figure 2A, B**. RT-qPCR was performed using common primers designed across the CAG intron upstream of the rescuing transgene. Values were normalised over TBP and expressed relative to S2R cells. Error bars represent the standard error of the mean (n=3).

To further assess the ability of SOX proteins to sustain ESC self-renewal, we attempted to expand transfected cell populations. PCR analysis confirmed deletion of the *Sox2* conditional allele (400bp) from expanded cell populations expressing SOX1, SOX2 or SOX3 (Figure 2 – Figure supplement 1). Furthermore, SoxB1-rescued (S1R and S3R) SKO ESCs retained OCT4 and NANOG expression, similar to SCKO and Sox2-rescued SKO (S2R) ESCs (Figure 2 – Figure supplement 2). The transgene expression levels in ESC populations rescued by Sox1, 2 or 3 were assessed by quantitative transcript analysis using common primers across the CAG intron. This demonstrated that Sox3 transgene mRNA was expressed at a higher level than the Sox1 or Sox2 transgenes (Figure 2D). These results confirm the findings of Niwa *et al.* in an independent cell line (Niwa *et al,* 2016).

### *Sox3* is dispensable for both naïve and primed pluripotency

Sox3 transcripts are present in ESCs at lower levels than Sox2 (Figure 3 – Figure supplement 1) and a previous report has suggested that Sox3 is dispensable for ESC selfrenewal (Rizzotti and Lovell-Badge, 2007). To directly assess this possibility, and as a first step to determining whether Sox3 is required in EpiSCs, CRISPR/Cas9 was used to delete the *Sox3* gene from male E14Tg2a ESCs (Hooper *et al,* 1987) using two guide RNAs (sgRNA1 and sgRNA2) (Figure 3A). After targeting, E14Tg2a ESCs were plated at low density and single clones were isolated. PCR genotyping using primers flanking the predicted Cas9 cut sites (Figure 3A) identified two clones (S3KO8 and S3KO35) in which *Sox3* had been deleted (Figure 3B). Replating at clonal density indicated that both clones retained an efficient self-renewal ability (Figure 3C). Moreover, both clones had unchanged levels of Nanog, Oct4 and Sox2 mRNAs (Figure 3D). These findings indicate that *Sox3* is dispensable for ESC self-renewal, confirming previously unpublished data (Rizzoti *et al,* 2004).

**Figure 3:**
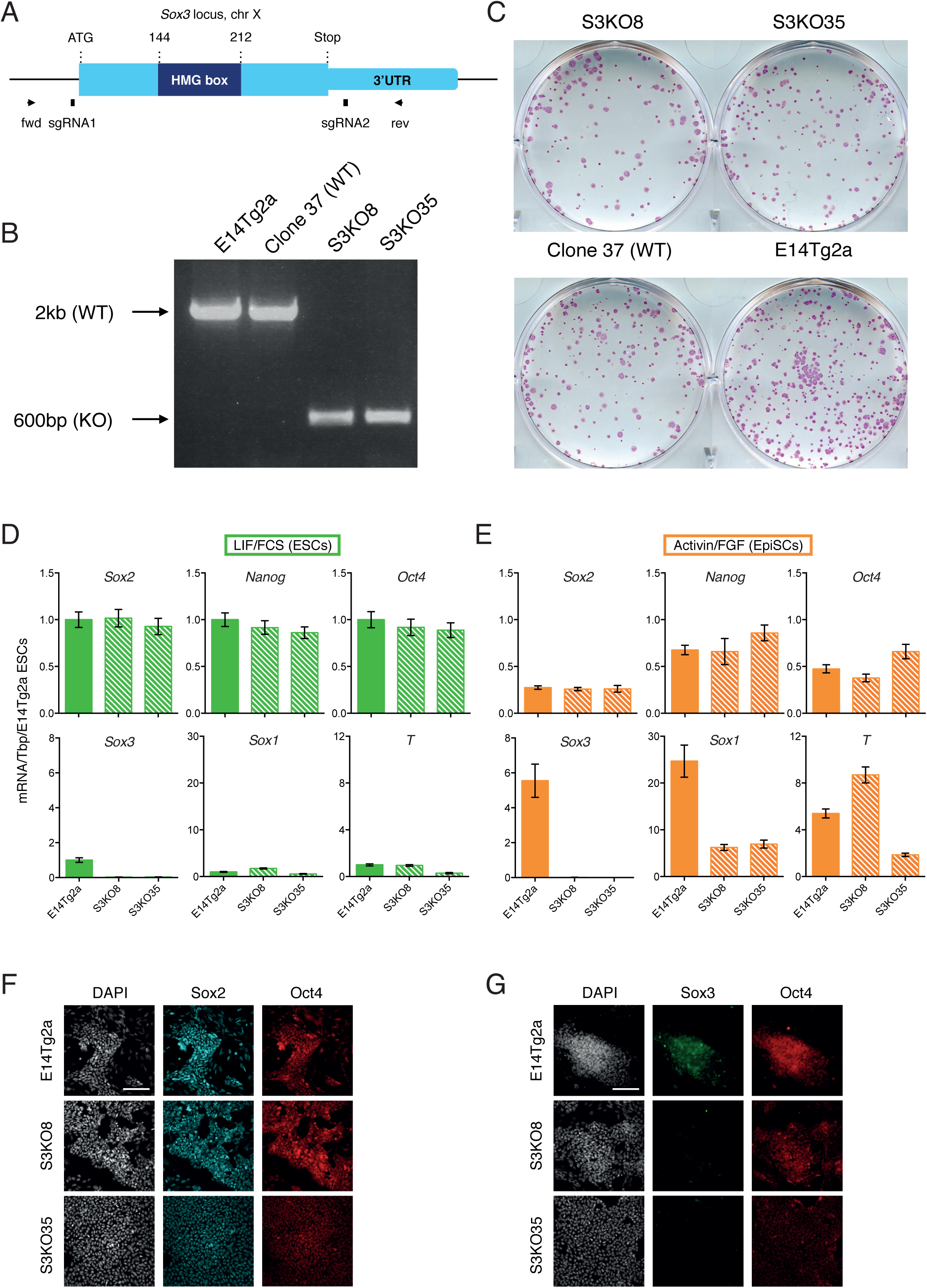
*Sox3* is dispensable for pluripotent cell maintenance. A. Schematic representation of the *Sox3* locus on the mouse X chromosome showing the CRISPR/Cas9 strategy for the generation and genotyping of *Sox3* knockout ESCs. The positions of sgRNA1 and sgRNA2 used for the deletion of the *Sox3* locus are indicated alongside the positions of PCR primers used for genotyping. B. PCR genotyping of parental ESCs (E14Tg2a), a wild-type clone (37) and two *Sox3* knockout clones (S3KO8, S3KO35) after CRISPR/Cas9 targeting of the *Sox3* locus. Band sizes for the wild-type (~2kb) and the targeted (~600bp) allele are shown. C. Alkaline phosphatase staining in *Sox3* knockout (S3KO8, S3KO35) and WT (Clone 37 and E14Tg2a) ESCs grown at clonal density for 7 days. D-E. Quantitative transcript analysis of the indicated transcripts in wild-type E14Tg2a and *Sox3* knockout (S3KO8, S3KO35) ESCs (D) and EpiSCs (E) derived by serial passaging (>10 passages) in Activin/FGF conditions. Error bars represent standard error of the mean (n=3). F. -G. Immunofluorescence analysis of SOX2 (F), SOX3 (G) and OCT4 proteins in E14Tg2a and *Sox3* knockout EpiSCs (S3KO8, S3KO35). Scale bar, 100μm.

Next, the requirement of SOX3 for primed pluripotency was determined by examining the ability of the above *Sox3* knockout ESC clones, together with parental E14Tg2a ESCs, to be converted to EpiSCs by serial passaging in Activin and FGF (Guo *et al,* 2009). Quantitative transcript analysis at passage 12 indicated similar levels of Sox2, Oct4 and Nanog mRNA expression in wild-type and *Sox3* knockout EpiSCs (3). In contrast, Sox1 mRNA levels were reduced in both *Sox3* knockout clones (Figure 3E) and T (Brachyury) was more variably expressed (Figure 3E). The ability of EpiSCs to self-renew in the absence of SOX3 was maintained over 25 passages without affecting SOX2 and OCT4 protein expression or EpiSC morphology (Figure 3F, G). These data demonstrate that both naïve and primed pluripotent cells can self-renew in the absence of Sox3.

### *Sox2* is dispensable for the maintenance of primed pluripotency

To investigate whether primed pluripotency can be maintained in the absence of SOX2, SCKO EpiSCs were derived by *in vitro* culture of SCKO ESCs (Guo *et al,* 2009). SCKO EpiSCs were then transfected with a plasmid encoding both Cre recombinase and tdTomato (Figure 4A), since the CAG CreERT^2^ transgene (Figure 2A) had been silenced during the EpiSC transition. After 12-24 hours, tdTomato-positive EpiSCs were sorted by FACS and replated at low cell density to allow expansion of single clones. PCR genotyping of expanded EpiSC clones revealed that the *loxP*-flanked *Sox2* allele had been excised generating *Sox2*^-/-^ (SKO) clones (Figure 4A). Immunoblot analysis confirmed that these SKO clones did not express SOX2 protein and that *Sox2^fl/-^* EpiSCs expressed SOX2 at 50% the level of *Sox2*^+/+^ EpiSCs (Figure 4B). Moreover, NANOG protein levels were similar to parental SCKO EpiSCs or wild-type E14Tg2a EpiSCs ( gure 4B). *Sox2*^-/-^ EpiSCs retained an undifferentiated morphology (Figure 4C) and immunostaining confirmed the absence of SOX2 and continued OCT4 expression compared to parental SCKO EpiSCs (Figure 4D). Quantitative transcript analysis of Oct4, Nanog and Sox2 mRNA levels, indicated that, while Sox2 mRNA was absent from *Sox2*^-/-^ EpiSCs, both Oct4 and Nanog mRNA levels were unaffected (Figure 4E). This suggests that the core pluripotency gene regulatory network remains active even in the absence of SOX2 and that, in contrast to the situation with naïve pluripotency, primed pluripotency is not critically dependent upon Sox2.

**Figure 4.**
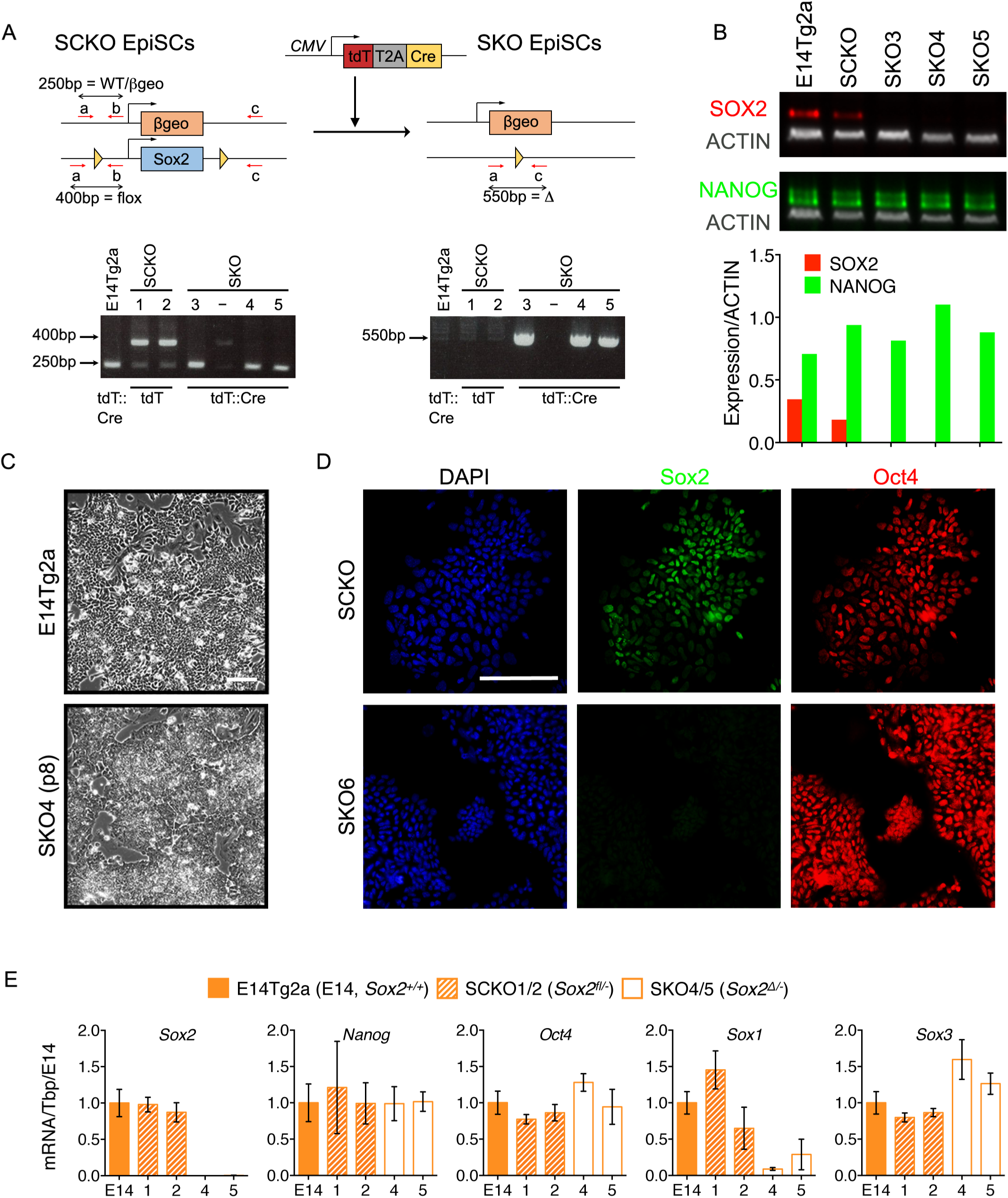
*Sox2* is dispensable for the maintenance of EpiSCs. A. Strategy for *Sox2* deletion in EpiSCs. *Sox2^fl/-^* (SCKO) EpiSCs were transfected with pCMV-tdTomato-T2A-Cre. After 12-24 hours cells were sorted for tdTomato expression and re-plated in the presence of ROCK inhibitor. Clones were picked and expanded before genotyping by PCR as indicated in Figure 1. SCKO EpiSCs were also transfected with pCMV-tdTomato (lacking Cre). Expanded clones were numbered as indicated. B. Immunoblot analysis of E14Tg2a, *Sox2^fl/-^* (SCKO) and *Sox2*^-/-^ (SKO) EpiSCs showing that SOX2 protein (red) is reduced in SCKO and absent in SKO EpiSCs. NANOG protein levels (green) were unaffected. Protein levels (normalized to the βactin level) are graphed below. C. Bright-field morphology of E14Tg2a and SKO4 EpiSCs after 2 weeks (8 passages) in culture. Scale bar, 100μm. D. Immunofluorescence analysis of EpiSCs for Sox2 and Oct4 showing the absence of Sox2 in SKO6 EpiSCs. Scale bar, 100μm. E Quantitative mRNA analysis of the indicated transcripts in E14Tg2a, *Sox2^fl/-^* (SCKO) and *Sox2*^-/-^ (SKO) EpiSCs. Clone numbers are indicated. Error bars represent the standard error of the mean (n=2 to 3).

The ability of SOXB1 proteins to substitute for one another in ESCs raised the hypothesis that functional redundancy between SOX2 and SOX3 may be responsible for the maintenance of pluripotency in primed EpiSCs. Expression of both Sox1 and Sox3 mRNAs was increased in EpiSCs compared to ESCs (Figure 1E). Examination of *Sox2*^-/-^ EpiSCs showed that while Sox1 mRNA expression was reduced, Sox3 mRNA levels were elevated compared to control cells (Figure 4E). This suggests that Sox3 expression might functionally compensate for the lack of Sox2 in *Sox2*^-/-^ EpiSCs, as is the case in *Sox2*-null E14.5 forebrain, where a 2-fold increase in Sox3 mRNA was detected (Miyagi *et al,* 2008). Alternatively, SOXB1 proteins may be irrelevant for EpiSC self-renewal. To distinguish between these possibilities we developed an approach to gene disruption using CRISPR/Cas9.

### Testing the functional effects of SOX ORF disruption using CRISPR/Cas9

To examine the functional dependence of cells on Sox genes, a CRISPR/Cas9 approach was developed and tested initially using *Sox2.* Insertions or deletions (indels) into the *Sox2* coding sequence were introduced immediately upstream of the sequence encoding for the SOX2 HMG domain (Figure 5 – Figure supplement 1A). Out of frame indels in this position are expected to abolish the DNA binding ability of any aberrant protein that might still be produced. The frequency and length of indels induced by CRISPR/Cas9-mediated targeting of the endogenous *Sox2* locus were investigated by TIDE (<u>T</u>racking of <u>I</u>ndels by <u>D</u>ecomposition) analysis (Brinkman *et al,* 2014). *Sox2^fl/-^* SCKO ESCs that constitutively express Sox1, Sox3 or GFP (generated in Figure 2B in the absence of tamoxifen) were analysed. If, as expected, Sox1 or Sox3 can rescue self-renewal, then such cells would carry both in-frame and out-of-frame (deleterious) indels (Figure 5 – Figure supplement 1B). In contrast, if ESC self-renewal relies on the remaining *Sox2* allele, as anticipated in the case of SCKO ESCs constitutively expressing GFP, then cells carrying deleterious indels should be eliminated from the population. In this case the only modifications present would be non-deleterious in-frame indels (Figure 5 – Figure supplement 1B). SCKO ESCs expressing Sox1, Sox3 or GFP were transfected with a plasmid encoding the sgRNA and eCas9. Cells were selected, genomic DNA isolated, PCR amplified, sequenced and analysed by TIDE. The population of SCKO ESCs expressing GFP contained *Sox2* loci with no out-of-frame indels detected (Figure 5 – Figure supplement 2). In contrast, SCKO ESCs constitutively expressing Sox1 or Sox3 tolerated out of frame, deleterious indels at *Sox2* (size +1, -10, -13) (Figure 5 – Figure supplement 2). These results show that indel analysis can be applied to study functional redundancy between SOXB1 group members in the maintenance of pluripotent cells.

### *Sox2* and *Sox3* are functionally redundant for the maintenance of EpiSCs

Having established the utility of CRISPR/Cas9-mediated indel induction for gene function analysis, a similar strategy was applied to *Sox2*^-/-^ EpiSCs to determine whether SOXB1 proteins operate in a functionally redundant way to maintain EpiSCs. Since *Sox2*^-/-^ EpiSCs showed an increase in Sox3 mRNA expression upon deletion of the second *Sox2* allele, we focussed on *Sox3,* which is X-linked and thus present in only one copy in these male EpiSCs. Two sgRNAs (sgRNA 1 and 2) were designed to independently target the *Sox3* ORF immediately upstream of the sequence encoding the HMG DNA binding domain and tested individually (Figure 5A). TIDE analysis (Figure 5B) showed that in control cells expressing endogenous Sox2 (E14Tg2a and SCKO) both in frame and out of frame indels within *Sox3* were detected using either sgRNA (Figure 5C). In contrast, two independent *Sox2*^-/-^ EpiSC lines (SKO1, SKO6) retained only in frame deletions that did not disrupt the SOX3 HMG box (Figure 5C). Together with earlier results on *Sox3^null^* EpiSCs (Figure 3), these data indicate that EpiSCs lacking either *Sox2* or *Sox3* alone can be maintained, while cells lacking both SOX2 and SOX3 proteins are lost. Therefore, EpiSC self-renewal requires SOXB1 function and this can be provided by the expression of either endogenous *Sox2* or *Sox3*.

**Figure 5.**
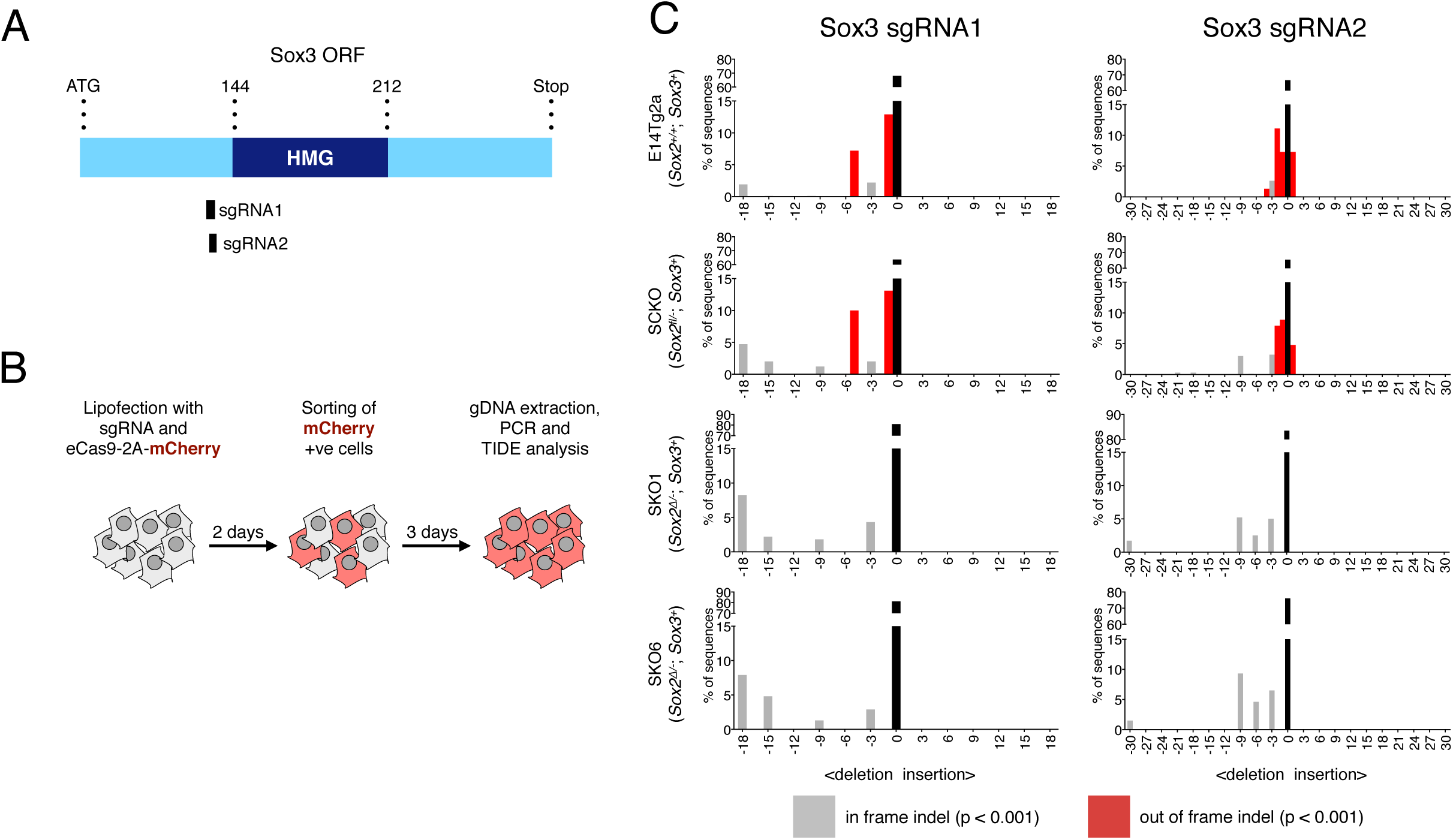
Sox3 indel analysis in *Sox2*^-/-^ EpiSCs. A. Schematic representation of the Sox3 coding sequence (ORF) with the HMG box highlighted in dark blue. The amino-acid positions of the HMG box relative to the start (ATG) codon of the ORF are shown. The positions of the sgRNAs 1 and 2 used are represented as black bars. B. Experimental design for Sox3 indel analysis in *Sox2*^-/-^ EpiSCs. C. Indel analysis performed using the TIDE tool (https://tide-calculator.nki.nl/) in two *Sox2*^-/-^ EpiSC lines (SKO1 and SKO6), in the parental *Sox2^fl/-^* (SCKO) EpiSCs and in control E14Tg2a EpiSCs using Sox3 sgRNAs 1 and 2. Histograms represent indel frequency and size. Black bars indicate the frequency of unmodified (WT) alleles; grey bars indicate significant in frame indels and red bars indicate significant out of frame indels (p<0.001). Not significant (n/s, p≥0.001) indels are not shown.

**Figure 6:**
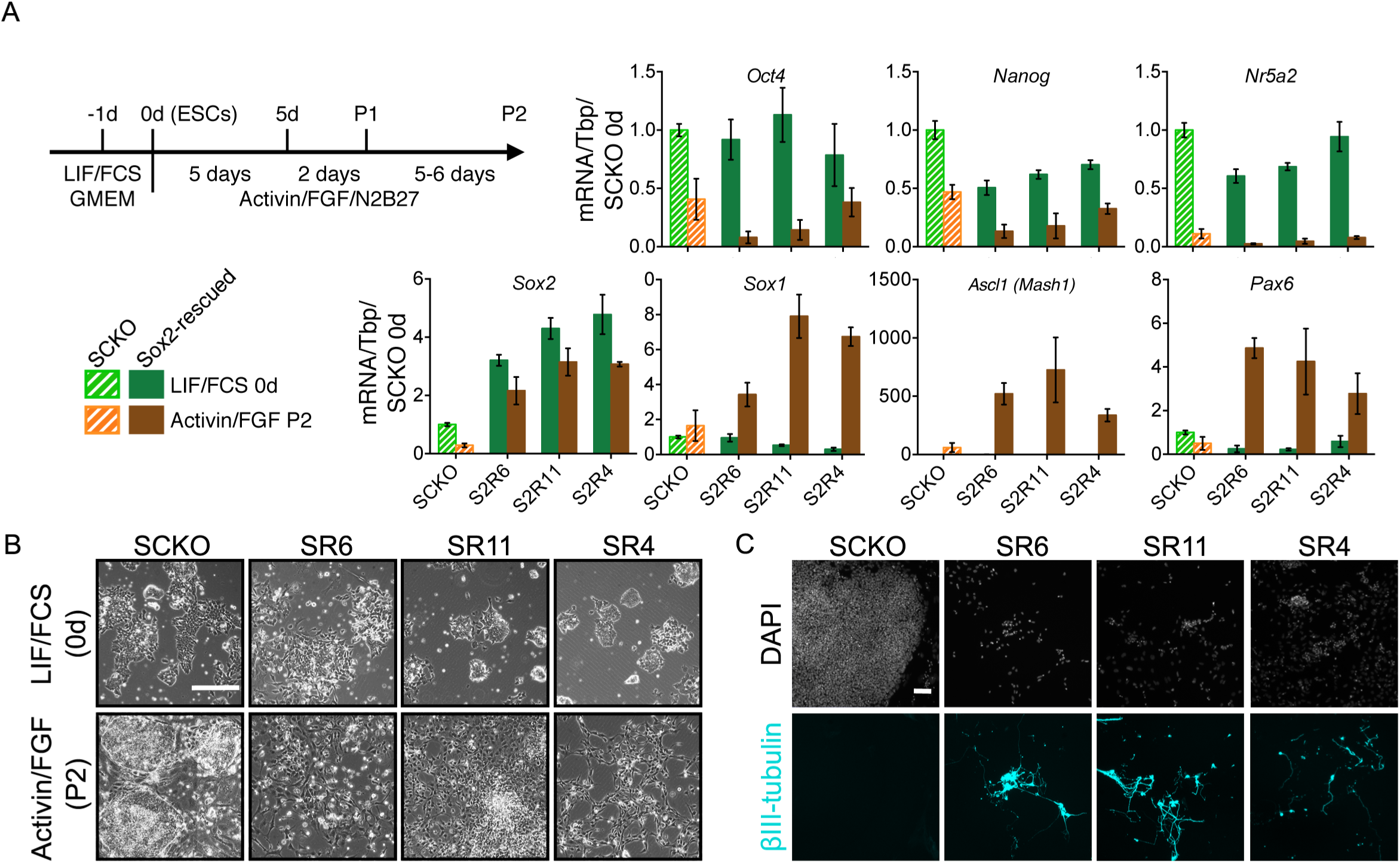
Increased SOXB1 levels skew ESC differentiation towards the neural lineage. A. Schematic diagram showing experimental plan. RT-qPCR analysis of the indicated mRNA levels in SCKO cells stably transfected and rescued with a Sox2 transgene (12-24h prior to a 24h treatment with 4OHT) (termed <u>S</u>ox<u>2</u>-<u>r</u>escued, S2R cells). Cells were grown in LIF/FCS conditions or differentiated in Activin/FGF conditions for 13 days. Transcript levels were normalized to TBP and plotted relative to SCKO ESCs. Error bars represent standard error of the mean (n=3 to 5). B. Bright field images of the indicated cells maintained in LIF/FCS or differentiated in Activin/FGF conditions (P2). Scale bar, 100μm. C. Immunofluorescence staining of the neural marker βIII-tubulin (cyan) in cells differentiated in Activin/FGF conditions (P2); DAPI represented in grey. Scale bar, 100μm.

### Modulating SOXB1 levels affects differentiation and can prevent capture of primed pluripotency

Since Sox2 and Sox3 transcript levels change during ESC to EpiSC differentiation, this raised the question of their importance for attainment of a primed pluripotent state. ESCs that delete *Sox2* differentiate to trophectoderm (Masui *et al,* 2007), while ESCs that continue expressing high SOX2 protein levels during differentiation are biased towards neural fates (Zhao *et al,* 2004). To assess the effect of increasing the SOXB1 concentration upon differentiation, we examined three clones overexpressing either Sox2 (Figure 6) or Sox3 (Figure 6 – Figure supplement 1), generated using the approach outlined in Figure 2A, When placed in an EpiSC differentiation protocol, both SOX2- and SOX3-overexpressing clones showed reduced expression of Oct4, Nanog and Nr5a2 transcripts as well as increased expression of Sox1, Mash1 and Pax6 transcripts (Figure 6A; Figure 6- figure supplement 1). Examination of SOX2-overexpressing clones in Activin/FGF showed neural-like cellular morphology (Figure 6B) and βIII-tubulin-positive axonal processes (Figure 6C). These findings indicate that while high SOXB1 levels are tolerated by ESCs, a decreased dosage of SOXB1 is essential for ESCs to transit effectively to an EpiSC state and avoid ectopic neural differentiation.

To further assess the effect of reducing the SOXB1 concentration upon ESC differentiation, ESCs lacking one *Sox2* allele were examined. Initially, we compared ESCs previously cultured in LIF/FCS or LIF/2i during neural differentiation induced by culture in N2B27 medium (Ying & Smith, 2003; Ying *et al,* 2003a). Our results indicate a more rapid induction of a Sox1-GFP reporter and Sox1 mRNA from LIF/FCS cultures than from LIF/2i cultures (Figure 7A). The Sox1-GFP kinetics from 2i/LIF cultures that we observe here agree with those reported (Marks *et al,* 2012), although this study noted a slower induction of Sox1-GFP from FCS/LIF cultures. However, as the timing of Sox1-GFP induction from FCS/LIF cultures in previous studies from the same group (Ying *et al,* 2003a) was consistent with the timings we report here, we initiated further differentiation experiments from LIF/FCS. Expression of pluripotency markers Nanog and Nr5a2 was decreased in the neural differentiation protocol, although with lower efficiency in *Sox2^fl/-^* ESCs than in control cells, while Fgf5 was induced (Figure 7B). Strikingly, Sox1, Mash1 and Pax6 mRNAs were induced in *Sox2*^+/+^ but not *sox2^fl/-^* cells (Figure 7B), and an increase in Sox3 was not sustained. These data indicate that a SOXB1 level above that present in *sox2^fl/-^* cells is required to enable ESCs to undergo neural differentiation *in vitro.* Furthermore, the expression of Fgf5 suggests that *sox2^fl/-^* cells attain some aspects of an early postimplantation identity (Figure 7B). This contrasts with the neural differentiation observed in *sox2*^+/-^ embryos, suggesting compensatory mechanisms other than SOXB1 redundancy exist in the embryo (Avilion *et al,* 2003; Rizzoti & Lovell-Badge, 2007; Favaro *et al,* 2009).

**Figure 7:**
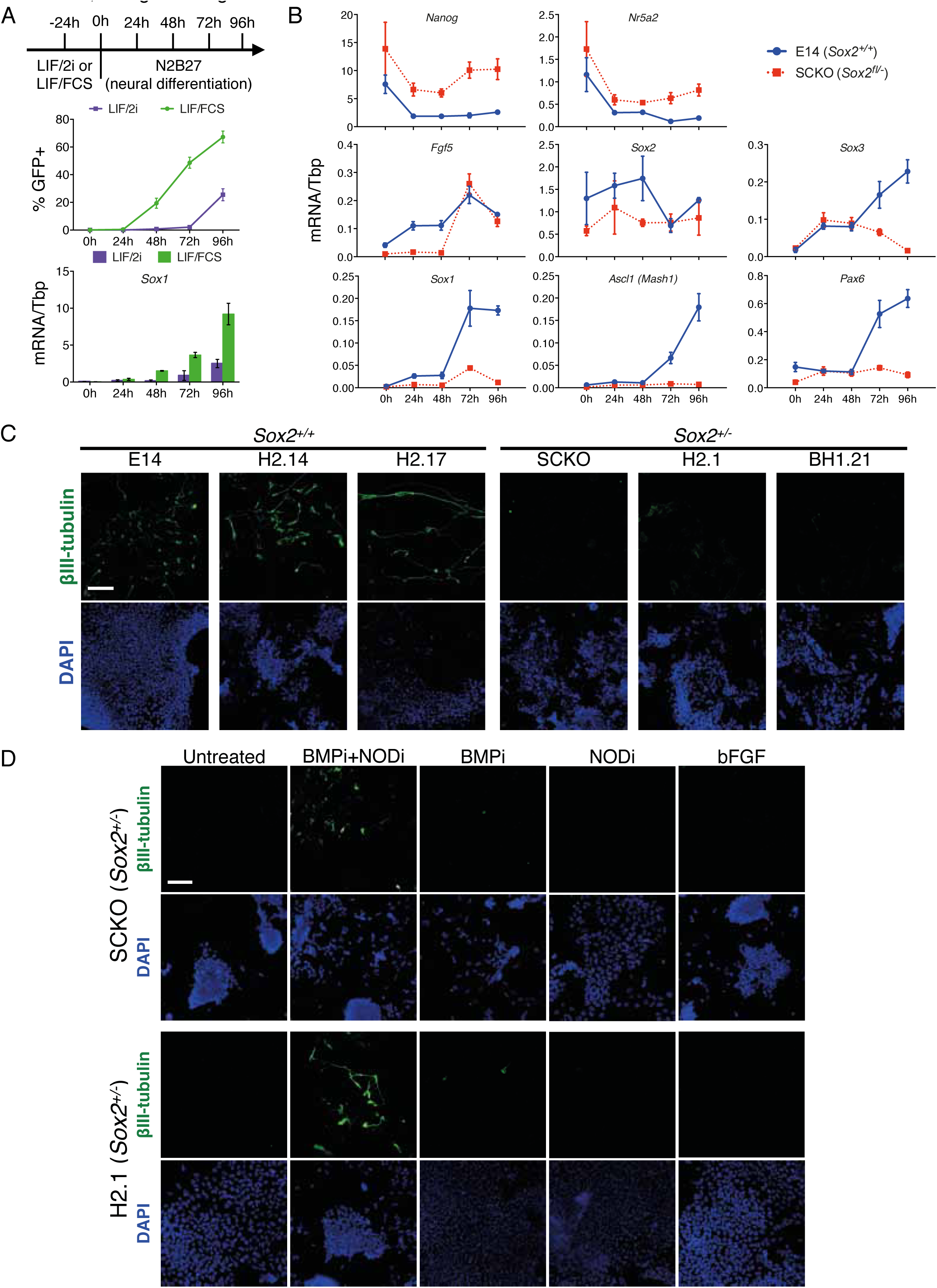
Decreased SOX2 levels prevent ESC differentiation into neurons. A. (top) Schematic diagram showing experimental plan for neural differentiation. (middle) Sox1-GFP (Aubert *et al,* 2003) expression in 46C cells cultured in LIF/FCS (green) or LIF/2i (purple) and for the indicated number of hours in N2B27 neural differentiation medium were assessed by flow cytometry. The positive (+) gate was set above the GFP expression level observed in 46C ESCs. The error bars indicate the standard error of the mean (n=4). (bottom) Quantitative mRNA analysis of *Sox1* mRNA level in 46C cells during neural differentiation. Error bars indicate the standard error of the mean (n=2). B. Quantitative mRNA analysis showing mRNA levels of the indicated transcripts in differentiating E14Tg2a (E14, *Sox2*^+/+^) and SCKO (*Sox2^fl/−^*) cells. Transcript levels were normalized to TBP. Error bars represent standard error of the mean (n=3). C. Immunofluorescence staining of the neural marker βIII-tubulin (green) in *Sox2*^+/+^ E14Tg2a (E14), H2.14 and H2.17 ESCs, and in *Sox2*^+/-^ SCKO, H2.1 and BH1.21 ESCs differentiated in N2B27 medium for 4 days; DAPI represented in blue. Scale bar, 100μm. D. Immunofluorescence staining of the neural marker βIII-tubulin (green) in *Sox2*^+/-^ SCKO and H2.1 ESCs differentiated in N2B27 medium for 4 days in the presence or in the absence of the LDN-193189 BMP inhibitor (BMPi), of the SB-431542 Nodal inhibitor (NODi) and of recombinant bFGF; DAPI represented in blue. Scale bar, 100μm.

To eliminate the possibility that the neural differentiation defect in SCKO cells resulted from a defect unrelated to *Sox2,* CRISPR/Cas9 was used to introduce indels in the *Sox2* ORF in E14Tg2a ESCs. Using independent sgRNAs, two *Sox2*^+/-^ ESC clones were isolated. Clone H2.1 carried an 8-bp deletion on one *Sox2* allele; clone BH1.21 carried a 10-bp deletion on one allele and a 3-bp deletion on the other allele (Figure 7 – Figure supplement 1A). Two additional clones in which the *Sox2* alleles were not modified (H2.14, H2.17) were used as controls. The -8 and -10 deletions cause frame-shifts introducing stop codons, while the -3bp deletion is likely to be functionally neutral as it occurs N-terminal to the HMG domain (Figure 7 – Figure supplement 1B). Placing *Sox2*^+/+^ ESCs (H2.14, H2.17 and E14Tg2a) in a neural differentiation protocol produced βIII-tubulin positive neurons. In contrast, no βIII-tubulin positive cells were detected from parallel treatments of H2.1, BH1.21 and SCKO *sox2*^+/-^ ESCs (Figure 7C). This establishes that the lack of a functional *Sox2* allele impairs effective neural differentiation of ESCs.

Neural differentiation of pluripotent cells is stimulated by FGF (Ying *et al,* 2003a; Ying & Smith, 2003) and inhibited by both BMP and Nodal/Activin (Ying *et al,* 2003b; Vallier *et al,* 2004, 2009; Guo *et al,* 2009). To determine whether perturbations in these pathways could overcome the neural differentiation defect of *Sox2*^+/-^ ESCs, H2.1 and SCKO cells were placed in the neural differentiation protocol supplemented with either recombinant bFGF or inhibitors of BMP or Nodal. Additional FGF, or inhibition of Nodal alone, was without effect, while BMP inhibition resulted in only a few βIII-tubulin-positive cells. However, simultaneous BMP and Nodal inhibition enabled *Sox2*^+/-^ ESCs to form βIII-tubulin-positive cells (Figure 7D). These results indicate that a reduction in SOX2 levels in ESCs enhances the response of cells to endogenous BMP and Nodal signalling, preventing effective neural differentiation.

To further investigate the effects of modulating SoxB1 gene dosage upon ESC differentiation, *Sox3^null^* ESCs were generated by CRISPR/Cas9 mediated gene deletion from heterozygous *sox2*^fl/-^ ESCs (SCKO). PCR genotyping identified two *sox2^fl/-^*; *Sox3^null^* ESC clones (#36 and #37) (Figure 8A). Quantitative transcript analysis showed that while *sox2^fl/-^* ESCs had an expected reduction in Sox2 mRNA and unchanged levels of Sox1 and Sox3 mRNAs compared to E14Tg2a ESCs, deletion of *Sox3* from *Sox2^fl/-^* ESCs increased Sox1 and surprisingly, also Sox2 mRNA to a level similar to that present in E14Tg2a ESCs (Figure 8B). These data suggest cross-regulatory interactions between SoxB1 members in which SOX3 protein represses *Sox1* and *Sox2,* either directly or indirectly, in ESCs (Figure 8B). However, as *Sox3* deletion in *Sox2*^+/+^ ESCs did not increase Sox1 and Sox2 mRNA levels (Figure 3D) this suggests that the repressive effect of SOX3 on *Sox2* is sensitive to the SOX2 protein concentration.

**Figure 8:**
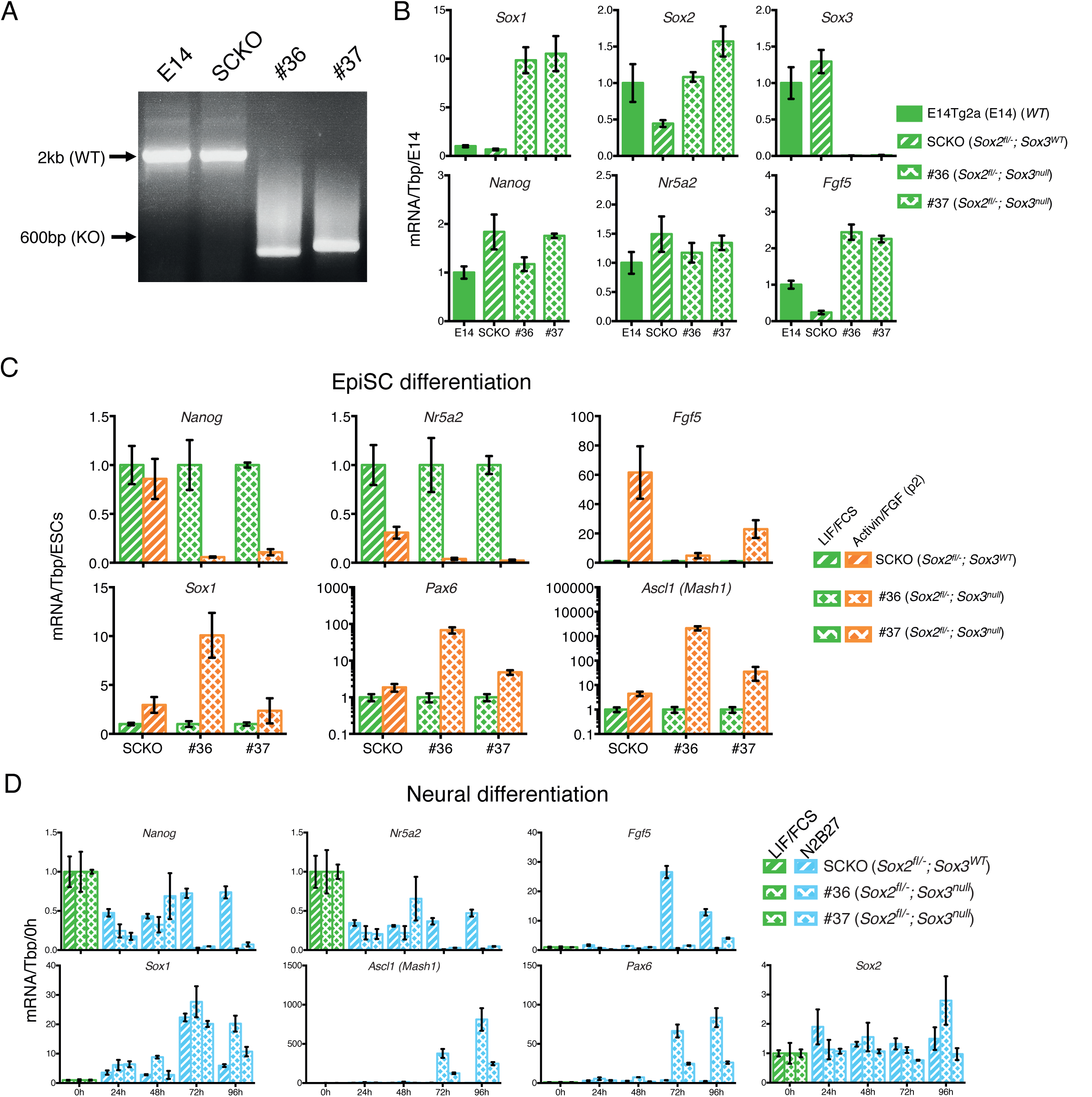
*Sox2/Sox3* requirements during EpiSCs differentiation. A. Genotyping analysis of the *Sox3* locus in E14Tg2a (E14, *Sox2*^+/+^), SCKO (*Sox2^fl/-^*) ESCs (≃2kb band) and in two *Sox3^null^; Sox2^fl/-^* ESC clones (#36 and #37) derived after deletion of the *Sox3* locus (≃600bp band). Gene targeting was performed following the strategy depicted in Figure 3A. B. Quantitative mRNA analysis showing mRNA levels of the indicated transcripts in E14Tg2a ESCs (E14, *Sox2*^+/+^), SCKO ESCs (*Sox2^fl/-^*) and in two *Sox3^null^; Sox2^fl/-^* ESC clones (#36 and #37). mRNA levels were normalised over TBP and plotted relative to E14Tg2a. Error bars indicate the standard error of the mean (n=3). C. Quantitative mRNA analysis showing mRNA levels of the indicated transcripts in SCKO cells (*Sox2^fl/-^*) and in two *Sox3^null^; Sox2^fl/-^* clones (#36 and #37) grown in LIF/FCS condition and in Activin/FGF conditions for 2 passages (P2). mRNA levels were normalised to TBP and plotted relative to ESC (LIF/FCS). Error bars indicate the standard error of the mean (n=3). D. Quantitative mRNA analysis showing mRNA levels of the indicated transcripts in SCKO cells (*Sox2^fl/-^*) and in two *Sox3^null^ Sox2^fl/-^* clones (#36 and #37) grown in LIF/FCS condition and in neural differentiation medium (N2B27) for the indicated number of hours. mRNA levels were normalised to TBP and plotted relative to ESC (LIF/FCS). Error bars indicate the standard error of the mean (n=3).

Next, the ability of *sox2^fl/-^; Sox3^null^* ESCs to transition to primed pluripotency was assessed by passaging in Activin/FGF. Whereas *sox2^fl/-^* EpiSCs could be successfully established and maintained (Figure 4), *sox2^fl/-^; Sox3^null^* cells displayed a differentiated morphology within 2 passages. Quantitative transcript analysis showed that in comparison to *sox2^fl/-^* cells, *sox2^fl/-^; Sox3^null^* cells induced less Fgf5 but more Mash1 and Pax6 (Figure 8C). These data suggest that while primed pluripotency can be maintained in the absence of either *Sox2* or *Sox3,* the transition from a naïve ESC state to a primed EpiSC state does not occur effectively when *Sox3* is absent and the *Sox2* gene dosage is halved. Notably, compared to *Sox2^fl/-^* ESCs, *Sox2^fl/-^; Sox3^null^* ESCs have the same Sox2 mRNA level as *Sox2*^+/+^ ESCs and Sox1 mRNA is increased >10-fold (Figure 8B). Such an increase in Sox1 mRNA expression is sufficient to enforce neural differentiation following LIF withdrawal (Zhao *et al,* 2004).

The increased levels of Sox1 and Sox2 mRNAs in LIF/FCS (Figure 8B), together with the increased levels of neural differentiation markers during EpiSC induction of *Sox2^fl/-^; Sox3^null^* ESCs (Figure 8C), prompted us to examine the behaviour of *Sox2^fl/-^; Sox3^null^* ESCs during neural differentiation. While *Sox2^fl/-^* ESCs did not effectively undergo neural differentiation (Figure 6), deletion of *Sox3* from *Sox2^fl/-^* ESCs was sufficient to rescue neural differentiation as judged by induction of Sox1, Mash1 and Pax6 mRNAs and of the βIII-tubulin protein (Figure 8D, Figure 8 – Figure supplement 1). This is likely a secondary consequence of the increase in Sox2/Sox1 mRNA expression in ESCs resulting from *Sox3* deletion (Figure 8B). The two *Sox2*^+/+^; *Sox3^null^* ESC clones (Figure 3A) were able to differentiate into neural cells with similar efficiency to parental cells as shown by induction of Sox1, Mash1 and Pax6 mRNAs (Figure 8 – Figure supplement 2) and appearance of βIII-tubulin-positive cells (Figure 8 – Figure supplement 1).

These data demonstrate that altering the SoxB1 genetic composition by deletion of *Sox3* and elimination of one functional *Sox2* allele induces a transcriptional de-regulation of the remaining *SoxB1* alleles. The increased mRNA levels of Sox1 and Sox2 are then sufficient to rescue neural differentiation of *Sox2^fl/-^* ESCs but also impair the ability of *Sox2^fl/-^; Sox3^null^* ESCs to be captured as primed EpiSCs.

## Discussion

Our examination of SoxB1 function in pluripotent cells extends previous findings that SOXB1 proteins can act redundantly in ESCs (Niwa *et al,* 2016) and during somatic cell reprogramming (Nakagawa *et al*, 2007) by showing that in EpiSCs, SOX2 and SOX3 proteins are functionally redundant within an altered PGRN. Functional redundancy implies that we can consider the total SOXB1 complement as a key factor in the outcomes selected by cells genetically engineered to express varying combinations of functional *SoxB1* alleles. In particular, a broad range of SOXB1 levels are tolerated by ESCs. However, upon exit from naïve pluripotency, low SOXB1 levels permit entry to the EpiSC state, while high levels enforce neural differentiation (Figure 9). We have also uncovered cross-regulatory relationships between Sox3 and the other SoxB1 members, that we hypothesise act to maintain an adequate SOXB1 level.

**Figure 9:**
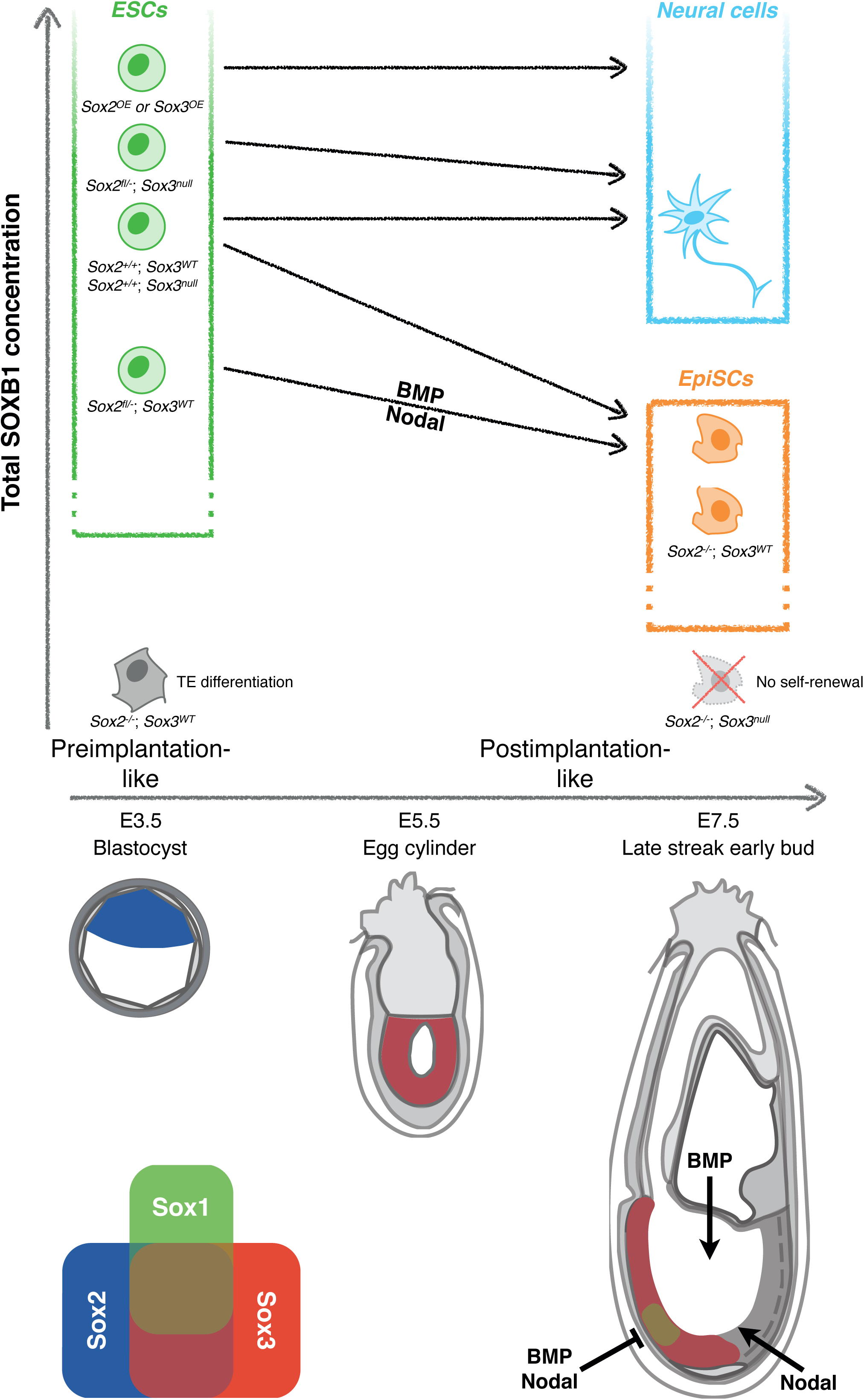
Dependence of cell fate potential of ESCs on the total SOXB1 concentration. The overall SOXB1 concentration inferred from SoxB1 transcript levels in different *SoxB1* mutant cell lines. Naïve ESCs self-renew in a wide range of SOXB1 concentrations. However, only ESCs with approximately wild-type SOXB1 levels can differentiate towards both primed EpiSCs and neural cells. ESCs with increased SOXB1 concentrations are poised towards neural differentiation, preventing their capture as primed EpiSCs in FGF/ActivinA. Decreased SOXB1 concentrations are insufficient to enable neural differentiation due to increased activity of neural antagonists (BMP and Nodal). A further reduction in SOXB1 is tolerated in the primed state due to SOX2/SOX3 functional redundancy but complete loss of the predominant SoxB1 forms is incompatible with self-renewal of both naïve and primed pluripotent cells. Depicted below are diagrams of pre- (E3.5) and postimplantation (E5.5 and E7.5) mouse embryos indicating the published expression patterns of SoxB1 mRNAs (Wood & Episkopou, 1999; Uchikawa *et al,* 2011; Avilion *et al,* 2003; Cajal *et al,* 2012), and the areas of BMP/Nodal signalling and inhibition during gastrulation (Constam & Robertson, 2000; Bachiller *et al,* 2000; Kinder *et al,* 2001; Levine *et al,* 2006; Pereira *et al,* 2012; Norris *et al,* 2002; Lawson *et al,* 1999; Perea-Gomez *et al,* 2002).

### SOXB1 function in primed pluripotent cells is provided by SOX2 and SOX3

In ESCs Sox2 mRNA is expressed at much higher levels in ESCs than either Sox1 or Sox3, and thus, in spite of redundancy, Sox2 can be considered to provide the dominant SOXB1 function in ESC pluripotency. However, even though *Sox2* is also the most abundant *Soxb1* transcript in EpiSCs, functional properties of the PGRN have changed. While Sox2 mRNA levels in EpiSCs are lower than in ESCs, *Sox3* levels are increased, and SoxB1 function is provided redundantly by Sox2 and Sox3. *Sox1* mRNA levels are also increased in these cells. The ability of Sox1 or Sox3 to substitute functionally for Sox2 in ESCs might suggest that Sox1 could also function redundantly with Sox2 to maintain EpiSCs. However, Sox1 may be less relevant to pluripotency as, unlike Sox2 and Sox3, which are widely expressed in the pluripotent postimplantation epiblast, Sox1 expression is uniquely associated with neural fate in the epiblast (Wood & Episkopou, 1999; Uchikawa *et al,* 2011; Cajal *et al,* 2012); Figure 9). Indeed, Sox1 expression in EpiSC populations is associated with neural committed cells (Tsakiridis *et al,* 2014).

### SOXB1 redundancy in vivo

In the seminal *Sox2* deletion study (Avilion *et al,* 2003) *Sox2^null^* embryos fail to develop a postimplantation epiblast. This was hypothesised to be due to the fact that neither Sox1 nor Sox3 were expressed sufficiently at the time of embryonic failure and therefore no redundantly acting SOXB1 protein could compensate for the SOX2 absence. Additional instances of potential SoxB1 redundancy have been reported *in vivo.* Replacement of an endogenous *Sox2* allele with a Sox1 ORF produced no phenotype, suggestive of functional interchangeability (Ekonomou *et al,* 2005). SOXB1 redundancy is likely to be evolutionarily conserved since in the chick, Sox2 and Sox3 both promote development of ectoderm and neurectoderm at gastrulation, although interestingly, in this case Sox3 expression occurs before Sox2 (Acloque *et al,* 2011). Moreover, while genetic knock-ins have shown that placement of Sox2 ORF at the *Sox3* locus can rescue pituitary and testes phenotypes caused by *Sox3* deletion (Adikusuma *et al,* 2017), the effects on pluripotent cells were not assessed. It is therefore interesting that while *Sox3^null^* mice can be viable, on a 129 genetic background they exhibit gastrulation-stage lethality (Rizzoti & Lovell-Badge, 2007; Adikusuma *et al*, 2017). This suggests that aspects of the regulation of SoxB1 expression that we show here to be important *in vitro* could also contribute to strain-specific differences in the timing or regulation of SOXB1 activity in vivo. Further studies will be required to establish the extent to which SOXB1 proteins can substitute genetically for one another in pluripotent cells *in vivo.*

### Low SoxB1 levels permit entry to the EpiSC state

Our results indicate that the total SoxB1 transcript level in ESCs needs to be reduced to enable entry into the EpiSC state rather than neural differentiation (Figure 9). *Sox2*^+/-^ ESCs placed in a neural differentiation protocol showed a reduced down-regulation of pluripotency markers (Nanog and Nr5a2) and failed to induce markers of neural differentiation. Surprisingly however, deletion of *Sox3* from SCKO cells to produce *Sox2^fl/-^; Sox3^null^* ESCs restored neural differentiation capacity (Figure 8D). Moreover, placement of *Sox2^fl/-^; Sox3^null^* ESCs in an EpiSC differentiation protocol resulted in a skewing of differentiation towards a neural identity as indicated by loss of Nanog, Nr5a2 and Fgf5 transcripts, an increase in mRNAs for neural differentiation markers (Figure 8C). This paradoxical behaviour may be explained by the fact that both Sox1 and Sox2 mRNAs increase upon elimination of *Sox3* from *Sox2^fl/-^* ESCs (Figure 8B). Interestingly, reciprocal repression of Sox3 by Sox2 also occurs in EpiSCs (Figure 4E). Unravelling the regulatory relationships between *SoxB1* genes is a relevant point for future studies.

### High SoxB1 levels enforce neural differentiation

Previous results have shown that elevating SOX1 or SOX2 expression in differentiating ESCs promotes neural differentiation (Zhao *et al,* 2004). In the present study, we showed that ESCs with either elevated SOX2 or SOX3 self-renew efficiently. However when these cells were placed in an EpiSC differentiation protocol, differentiation was skewed towards a neural identity despite the presence of FGF and neural-antagonising Activin signals (Vallier *et al,* 2004, 2009). This indicates that enforced expression of SOXB1 proteins overrides the signalling system that captures primed pluripotent cells *in vitro.*

### Parallels between SOXB1 function during in vitro and in vivo neurogenesis

The dynamics of ESC differentiation *in vitro* towards neural and EpiSC states show interesting parallels with early development *in vivo* (Figure 9). While ESCs and preimplantation embryos express only Sox2, early post-implantation embryos, and differentiating ESCs, express Sox2 and Sox3 but not Sox1. At gastrulation stages, which are transcriptomically similar to EpiSCs (Kojima *et al,* 2014), Sox2 and Sox3 are expressed widely in the epiblast. As noted above, Sox1 expression is restricted to a subdomain within the Sox2/Sox3-positive epiblast that overlaps extensively with the prospective brain (Wood & Episkopou, 1999; Cajal *et al,* 2012), where early neural differentiation occurs *in vivo.* Thus the region expressing all three SOXB1 members might undergo neural induction as a consequence of expressing high levels of SOXB1 proteins, while the rest of the epiblast experiences lower SOXB1 levels, enabling pluripotency to extend through gastrulation.

### An interplay between SOXB1 function and anti-neural signals

Inhibition of both BMP and Nodal signalling rescues the neural differentiation ability of *Sox2*^+/-^ ESCs. These results suggest that *Sox2*^+/-^ ESCs can initiate exit from naïve pluripotency but cannot complete neural differentiation due to enhanced responses of cells to endogenous anti-neuralising signalling by BMP and Nodal. The viability of *Sox2*^+/-^ mice (Avilion *et al,* 2003; Rizzoti & Lovell-Badge, 2007; Favaro *et al,* 2009) indicates that the *in vivo* environment is able to overcome these anti-neural signals. In wild-type embryos, the prospective brain is shielded from Nodal and BMP signalling by secreted inhibitors of these pathways, including Cer1, Lefty1/2 (Perea-Gomez *et al,* 2002), Chrd and Nog (Bachiller *et al,* 2000). Removal of either Nodal or BMP inhibition leads to absence of anterior neural tissue. Interestingly, the cells that express these inhibitors (including the anterior visceral endoderm and node) do not express SOXB1 proteins and are therefore likely to be functionally unaltered in *Sox2*^+/-^ embryos. The observation that the reduced SOXB1 concentration in *Sox2*^+/-^ pluripotent cells is compatible with neural differentiation, provided that endogenous anti-neuralising signals are blocked indicates that to fully understand how the choice between neural and primed pluripotency is made, it will be necessary to elucidate how the signalling environment connects to the SOXB1-driven transcriptional programme.

## Materials and Methods

### Cell culture

For a complete list of cell lines, their name, their genotypes and their original characterisation, see Supplementary File 2. All the cell lines used in this study were regularly tested for contaminations and were mycoplasma negative.

ESCs grown in LIF/FCS conditions were cultured on dishes coated with 1% gelatin (Sigma) and in GMEM medium (Sigma) supplemented with 1x non-essential aminoacids (Life Technologies), 1mM sodium pyruvate (Life Technologies), 2mM glutamine (Life Technologies), 100U/ml human LIF (Sigma), 10% ESC-grade FCS (APS) and 100μM β-mercaptoethanol (Life Technologies). G418 (200μg/mL, Sigma) was supplemented to maintain SCKO ESCs.

To adapt ESCs into LIF/2i/N2B27 (Ying *et al*, 2008) (LIF/2i) or LIF/BMP4/N2B27 (Ying *et al,* 2003b) (LIF/BMP) conditions, ESCs were replated in LIF/FCS on gelatin-coated plates for 24 hours before changing the culture media to N2B27 medium (Ying & Smith, 2003) supplemented with 100U/ml LIF, 1μM PD0325901 and 3μM CHIR99021 (Stemgent) (LIF/2i) or 10ng/ml BMP4 (Life Technologies) (LIF/BMP). Cells were passaged for at least 4 passages before using for analysis.

EpiSCs were derived from LIF/FCS-cultured ESCs as described previously (Guo *et al,* 2009; Osorno *et al,* 2012) and cultured on dishes pre-coated with 7.5μg/ml fibronectin without feeders in N2B27 medium (Life Technologies) ActivinA (20ng/ml, Peprotech), bFGF (10ng/ml, Peprotech). In brief, ESCs were plated at a density of 3×10^3^ cells/cm2; the equivalent of 30×10^3^ cells per well of a 6 well plate. Medium was changed to Activin/FGF conditions 24 hours after replating. Cells were passaged after 4-5 days in a 1:20 dilution for the first 8-10 passages. Stable EpiSC lines were propagated by passaging in a 1:10-1:20 dilution.

For neural differentiation, ESCs were cultured with minor modifications as previously described (Ying *et al,* 2003a; Ying & Smith, 2003). Briefly, ESCs were replated in LIF/FCS on gelatin-coated plates for 24 hours before changing the culture media to N2B27 medium (Life Technologies) only and allowed to grow for the indicated amount of time. When indicated, cells were differentiated in the presence of bFGF (10ng/ml, Peprotech), LDN-193189 (100nM, Stemgent) and/or SB-431542 (10μM, Calbiochem).

### *Sox2* deletion in ESCs and EpiSCs

Sox2 deletion in SCKO ESCs was performed similarly to previously described (Gagliardi *et al,* 2013; Favaro *et al,* 2009). In brief, 10^7^ SCKO ESCs grown in LIF/FCS conditions were transfected using Lipofectamine 3000 (Life Technologies) with 6-15μg of transgene-expressing plasmid the indicated test cDNA before replated in LIF/FCS at a density of 1.5×10^6^ per 10cm dish cultured overnight in the presence or in the absence of 4-hydroxytamoxifen (1μM, Sigma). Medium was changed 12 to 24 hours later to LIF/FCS medium supplemented with Hygromycin B (150μg/mL, Roche). Transfected cells were cultured for 8 to 10 days before they were stained for alkaline phosphatase (AP) activity (Sigma). Stained colonies were scored based on the presence of AP-positive cells within the colony. In parallel, unstained populations were expanded to generate cell lines before genotyping by PCR.

SCKO EpiSCs were transfected with ptdTomato-T2A-Cre by lipofection (Lipofectamine 2000, Life Technologies). After 12-24 hours cells were sorted for tdTomato expression and replated in the presence of ROCK inhibitor (Y-27632, Calbiochem) for 24 hours before medium was changed to remove ROCK inhibitor. Clones were expanded before genotyping by PCR.

### CRISPR/Cas9 deletion of the *Sox3* gene

Two sgRNAs were designed upstream of the ATG (sgRNA1) and downstream of the stop codon (sgRNA2) using an online CRISPR Design Tool (http://crispr.mit.edu/) and subsequently cloned in the *BbsI*-linearised pSpCas9(BB)-2A-GFP plasmid (Addgene 48138) as previously described (Ran *et al,* 2013). sgRNA sequences used in this study are listed in Suppl. Table 1. 10^6^ E14Tg2a ESCs were co-transfected with 1μg of sgRNA1-encoding plasmid and 1μg of sgRNA2-encoding plasmid using Lipofectamine 3000 (Life Technologies) following the manufacturer’s instructions. 24 hours after transfection, GFP-positive cells were sorted by FACS and replated at clonal density to allow the isolation of single colonies. After 8-10 days, single ESC colonies were expanded and the *Sox3* locus was genotyped by PCR.

### Indel induction and TIDE analysis

SgRNAs targeting the *Sox2* or *Sox3* ORF immediately upstream of the sequence encoding for the HMG box were designed using an online CRISPR Design Tool (http://crispr.mit.edu/). The Sox2 sgRNA was subsequently cloned in the *BbsI*-linearised pSpCas9(BB)-2A-Puro (PX459) V2.0 plasmid (Addgene 62988) as previously described (Ran *et al,* 2013). The resulting plasmid, or the empty vector control, were then transfected using Lipofectamine 3000 (Life Technologies) into SCKO ESCs constitutively expressing either SOX1, SOX3 or GFP. After 24 hours, ESCs were selected with a high concentration of 1.5μg/ml of puromycin for 24 hours to enrich for transfected cells, and then expanded for 72 hours. For *Sox3* indel analysis, two individual sgRNAs were cloned in *BbsI*-linearised pSpCas9(BB)-2A-mCherry that was obtained by fusing a 2A-mCherry cassette to the Cas9 CDS of the eSpCas9(1.1) plasmid (Addgene 71814). 10^6^ E14Tg2a (*Sox2*^+/+^), SCKO (*Sox2^fl/-^*), SKO1 (*Sox2*^-/-^) and SKO6 (*Sox2*^-/-^) EpiSCs were then transfected with 1μg of sgRNA-containing plasmids or empty empty vector using Lipofectamine 3000 (Life Technologies). 48 hours after transfection, mCherry-positive cells were FACS sorted, replated and expanded for 72 hours. Genomic DNA (gDNA) was extracted using the DNeasy Blood & Tissue kit (Qiagen) following the manufacturer’s instructions. gDNA was PCR amplified using the Q5 HotStart Polymerase (NEB) and primers flanking either the Sox2 sgRNA or Sox3 sgRNAs recognition sites. Primers and sgRNAs used in this study are listed in Supplementary File 3. PCR amplicons were purified and submitted for Sanger sequencing using the same forward primer that they had been generated with. Sanger sequencing electropherograms were then submitted for indel analysis with the TIDE tool (https://tide-calculator.nki.nl/, (Brinkman *et al,* 2014)) using amplicons obtained from cells transfected with empty vector as reference sequences. Indel analysis was performed using the default TIDE settings and a window of 27bp for SCKO ESCs and Sox2 indel induction, and a window of 30bp for EpiSCs and Sox3 indel induction. Indels with pvalue > 0.001 were scored as statistically significant.

### PCR genotyping

Genomic DNA (gDNA) was extracted from cells using DNeasy kits (Qiagen) according to the manufacturer’s recommended protocol. 100ng of gDNA was used per PCR reaction. Primers used in this study are listed in Supplementary File 3.

### Plasmid constructs

A mouse genomic BAC containing 5kb on either side of the *Sox2* stop codon (Source bioscience bMQ314D22) was shredded by pSC101-BAD-gbaA mediated recombineering using the pACYC177 backbone comprising the p15-origin and β-lactamase gene to produce pSox2-10kb. To construct the Sox2-T2A-H2B-tdTomato-IRES-Neo cassette, 89bp and 168bp immediately upstream and downstream of, and excluding the *Sox2* stop codon were PCR amplified to serve as homology arms for recombineering. The 89bp upstream fragment has a 5’ *Xho* I site and a GSG-T2A sequence at the 3’ end containing an in-frame *Fse* I site that preserves the T2A Gly-Pro residues and is followed by *Not* I, *Pac* I, *Asc* I, *San* DI, *Bam* HI, *Afl* II, *Nhe* I, *Cla* I and *Xho* I sites. The sequence of this fusion is shown below:

GGCTCCGGAGAGGGCAGAGGAAGTCTGCTAACATGCGGTGACGTCGAGGAGAATCCTGGGCCGGCCGCGGCCGCTTAATTAAGGCGCGCCGGGACCCGGATCCGCTTAAGGCTAGCATCGATTCTCGAG

The 89bp fragment was cloned into a PCR amplified, pUC19-derived 2kb minimal vector comprising β-lactamase gene and Col E1 origin with 150nt flanking sequences and a single *Xho* I site (pL). Individual features were PCR-amplified, flanked with unique sites and cloned as follows: H2B-tdTomato was cloned in-frame between *Fse* I and *Not* I, using TAA stop codon from *Pac* I; Gtx-IRES was cloned between *Pac* I and *Asc* I; neomycin phosphotransferase (Npt) was cloned between *San* DI and *Bam* HI, using the TAA stop codon from *Afl* II. The 168 bp downstream fragment is flanked with *Nhe* I at the 5’ side and *Cla* I at the 3’ end and was cloned between *Nhe* I and *Cla* I (pL-Sx-5HTiN3).

To insert a selection cassette for use in recombineering and targeting, a linker was made by annealing the following two oligonucleotides:

TCGAGCTTAAGGTCGACAGATCTCGATCGGCTAGCC

TCGAGGCTAGCCGATCGAGATCTGTCGACCTTAAGC

The linker was cloned into the *Xho* I site of a pTOPO-BluntII-derived, Zeocin-resistant version of pL (pZ-Linker). A 3.6kb *Bam* HI fragment containing PGK-EM7-Npt-pA and MC1-HSVtk-pA flanked by FRT sites (FNF) was subcloned from pBS-M179 (a kind gift from Dr. Andrew Smith) into the *Bgl* II site of pZ-Linker (pZ-FNF), destroying the *Bam* HI and *Bgl* II sites. This places an *Afl* II site on one side and *Nhe* I site on the other side of the FNF cassette and these are used to subclone into pL-Sx-5HTiN3 to make pL-Sx-5HTiNFNF3.

To make the targeting vector, a 7.1kb *Xho* I-*Cla* I fragment from pL-Sx-5HTiNFNF3 was transfected into *E. coli* containing pSox2-10kb and pSC101-BAD-gbaA to replace the *Sox2* stop codon by recombineering. Successful kanamycin-resistant recombinants carrying a 20kb targeting vector (pSox2AHTiN-FNF-10kb) were amplified at 37°C to restrict pSC101-BAD-gbaA replication and identified by diagnostic *Bam* HI digest.

The targeting construct was linearised, electroporated into E14Tg2a ESCs and G418 resistant colonies expanded and genotyped by southern blot analysis as described below. The FRT-flanked cassette was removed by transiently transfecting pPGK-FlpO into verified E14<u>T</u>G2a-derived <u>S</u>ox2-td<u>T</u>omato ESC clone 18 (TST18).

Overexpression constructs were generated by cloning open reading frames (ORFs) of indicated Sox genes, preceeded upstream by the Kozak consensus sequence (GCCGCCACC), into pPyCAG-*IRES*-Hyg vector between the *Xho*I and *Not*I sites (Chambers *et al,* 2003).

### Southern blot analysis

40μg of genomic DNA were digested with *Eco* RI (5’ probe analysis) or with *Hind* III (3’ and internal probe analysis) and assessed by Southern blot analysis using probes synthesised by PCR from the oligonucleotides indicated in Supplementary File 3.

### RNA level quantification

Total RNA was purified from cells using RNeasy mini kits (Qiagen) following the manufacturer’s protocol. cDNA was prepared from 1μg of total RNA using SuperScript III reverse transcription kits (Life Technologies) according to the recommended protocol and the final cDNA solution was diluted 1:10 prior to use. 2μl of cDNA solution was used per reaction with the Takyon Sybr Assay (Eurogentec). Primers used in this study are listed in Supplementary File 3.

### Microarray gene expression analysis

Total RNA (127ng/sample) from three independently replicated experiments was converted into biotin-labelled cRNA using the Illumina TotalPrep RNA amplification kit (Ambion). Microarray hybridization reactions were performed on a Mouse WG-6v2 BeadChip (Illumina). Raw data were normalised in R using the beadarray (Dunning *et al,* 2007), limma (Smyth, 2005) and sva (Leek & Storey, 2007) packages from the Bioconductor suite (Gentleman *et al,* 2004). Briefly, low-quality probes were removed from the input and data was then quantile-normalized. ComBat was used to account for batch effects (Leek & Storey, 2007) between microarrays run at different dates. Differential expression in the log2-transformed data was assessed with the limma algorithms (Smyth, 2005). Probes were considered differentially expressed if they showed a FDR-adjusted p-value of ≤ 0.1 and an absolute log_2_ fold change ≥ log2(1.5). Primary analysis results were uploaded to GeneProf (Halbritter *et al,* 2011) and mapped to Ensembl-based reference genes, collapsing multiple probes for the same gene by picking the most responsive probe (i.e., the probe with the highest absolute fold change across all pair-wise comparisons). Data are available in Supplementary File 1. Raw and processed microarray data have been submitted to GEO and are available to reviewers under accession code GSE99185.

### Immunofluorescence staining

Cells were fixed with 4% PFA (10 mins, RT) and permeabilised using PBS/0.1% (v/v) Triton-X100 (PBSTr) for 10 mins, before quenching with 0.5M Glycine/PBSTr (15 mins). Non-specific antigens were blocked using 3% (v/v) donkey serum/1% (v/v) BSA/PBSTr (1 hour, RT) before incubating with primary antibody in blocking solution (4°C, overnight). Cells were washed with PBSTr before incubating with donkey-raised secondary antibodies conjugated with Alexa-488, -568 or -647 in blocking solution (1-2 hours, RT). DAPI (1-2μg/mL, Molecular Probes) in PBS was added to cells for at least 30 mins before imaged using the Olympus IX51 inverted fluorescent microscope. Primary antibodies used were: α-SOX2 (Abcam ab92494; 1:400), α-NANOG (eBioscience 14-5761, 1:500) and α-OCT4 (Santa Cruz sc8628, 1:400), α-SOX3 antibody (ab42471, 1:400), α-βIII-tubulin (Covant, MMS-435P, 1:1500).

### Immunoblotting analysis

Cells were lysed with lysis buffer comprising 50mM Tris pH 8.0, 150mM NaCl supplemented with fresh 0.5% NP-40, 0.5mM DTT, 1× protease inhibitors cocktail (Roche) and 1.3μl of Benzonase (Novagen) (1 hour, 4°C). Concentrations of total protein extracts were quantified with an adaptation of the Bradford assay (Bio-Rad) according to the manufacturer’s recommended protocol. Samples were prepared by boiling 40μg of total protein extract with Laemmli buffer (1:1 mix) (BioRad) in a total volume of 25μl for 5 mins before chilling on ice (5 mins). Samples were analysed using Bolt™ 10% Bis-Tris+ SDS-PAGE (Novex) and electroblotted onto 0.2μm pore Whatman-Protran nitrocellulose membranes (Capitol Scientific) in transfer buffer comprising 25mM Tris/0.21M glycine/20% methanol. Membranes were blocked using 5% (w/v) low-fat milk in 0.01% (v/v) Tween-20/PBS (PBSTw) (1 hour, RT) before incubating with primary antibody in blocking solution (overnight, 4°C). Membranes were washed with PBSTw before incubating with donkey-raised secondary antibodies conjugated with IRDye 800CW (LI-COR 926-32213) and HRP-conjugated α-βActin (Abcam ab20272, 1:10000) antibody in blocking solution (1 hour, RT). HRP-staining was developed using a Super-signal West Pico kit (Pierce) (5 mins, RT) before imaging the membranes using LI-COR Odyssey Fc imager. The primary antibodies used were: α-Sox2 (Abcam ab92494; 1:1000), α-Sox3 antibody (ab42471, 1:1000) and α-Nanog (Bethyl A300-397A, 1:2000).

### Embryo manipulation

Mice were maintained on a 12h light/dark cycle. All animals were maintained and treated in accordance with guidance from the UK Home Office. Embryonic day (E)0.5 was designated as noon on the day of finding a vaginal plug. Morula aggregations, blastocyst injections and embryo transfer were performed using standard procedures. Chimeric, gastrulation-stage embryos were collected at E7.5 and imaged using an inverted fluorescence microscopy (Olympus IX51). Chimeric E9.5 embryos were fixed with 4% PFA at 4°C for 3h and cryosectioned as described before (Wymeersch *et al,* 2016). Sections were stained for GFP as described above.

## Acknowledgements

We thank Emily Erickson and Christine Watson (Cambridge) for suggesting the use of TIDE analysis; CRM Animal House, Microscopy Facility and FACS Facility staff for assistance; Donal O’Carroll for comments on the manuscript. This research is supported by grants from the Medical Research Council (IC and VW) and the Biotechnological and Biological Sciences Research Council of the United Kingdom (IC) and by a Japan Partnering Award from the Biotechnological and Biological Sciences Research Council (IC). FCKW was supported by an MRC studentship and a Centenary Award; EM received an award from the International Association for the Exchange of Students for Technical Experience.

---

**Figure 1 – Figure supplement 1.**
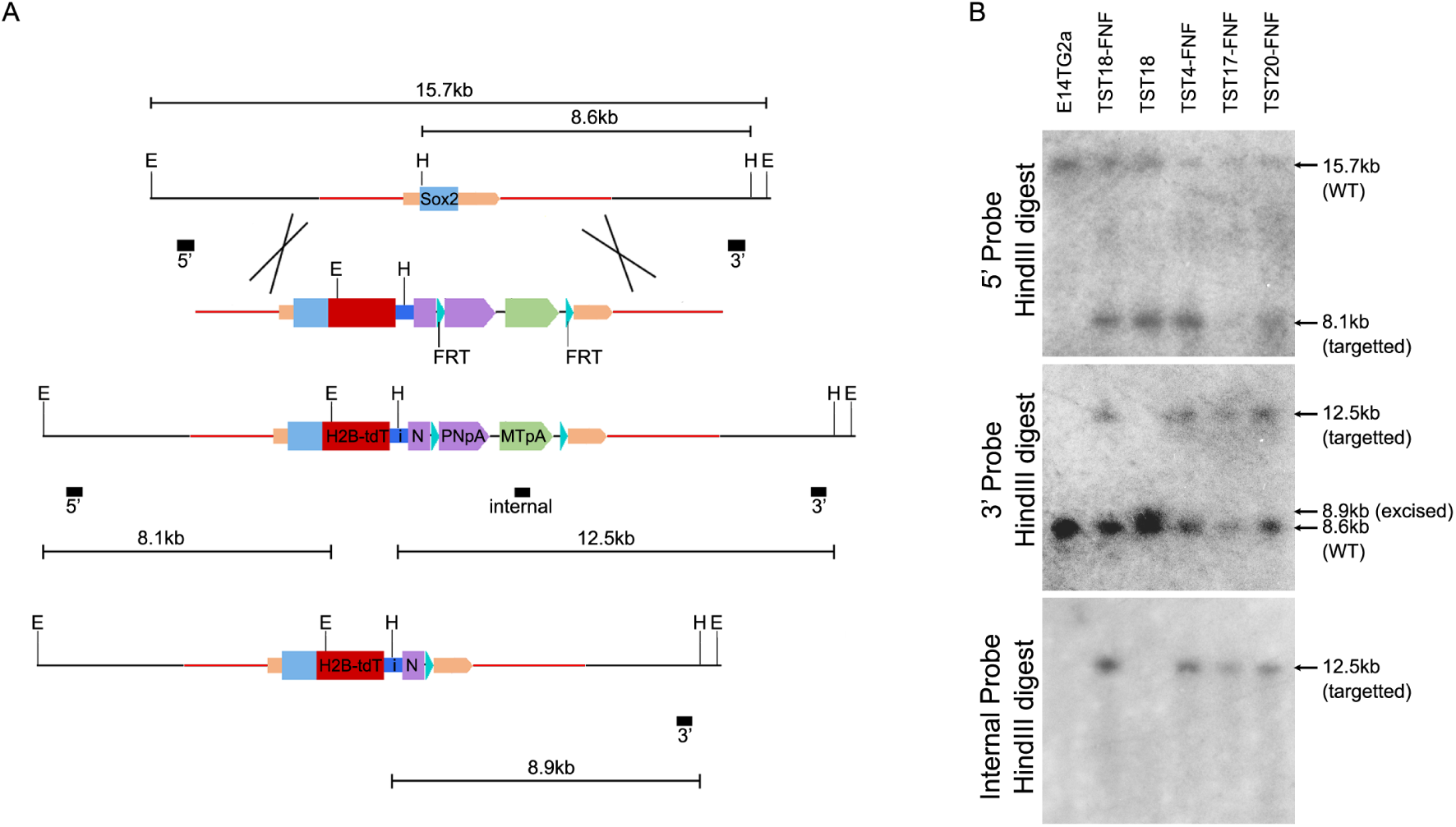
A. Targeting T2A-H2b-tdTomato to the *Sox2* locus stop codon yields the Sox2::HT fluorescent reporter allele. The Sox2 ORF is represented by a blue box and the Sox2 UTRs by beige boxes; homology arms are shown (red line). The Sox2::HT ORF is followed by GtxIRES-Neo and an FRT-flanked PGK-Neo-pA (PNpA)/MC1-TK-pA (MTpA) selection cassette. The FRT sites (cyan triangles), EcoRI sites (E), HindIII sites (H) and probes (black boxes) used for Southern analysis are shown. Sizes of relevant fragments produced before and after targeting and excision of the FRT-flanked cassette are indicated. B. Southern analysis of TST cell lines expanded after G418 selection. FNF denotes lines prior to excision of the FRT-flanked cassette.

**Figure 1 – Figure supplement 2.**
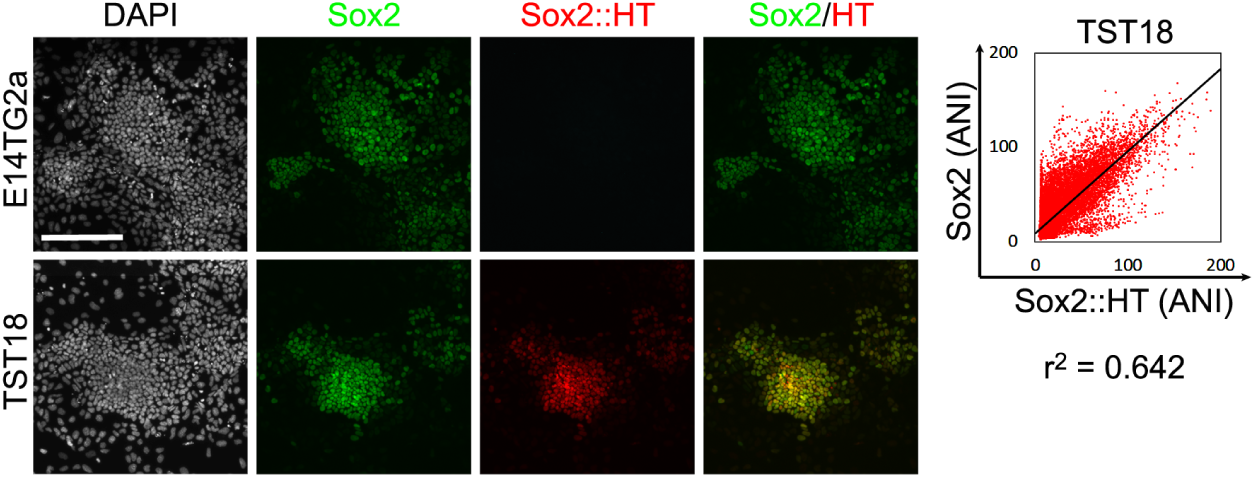
Imaging of E14Tg2a and TST (clone 18) ESCs for Sox2 by immunofluorescence (green) and of Sox2::HT fluorescence (red); DAPI is grey. Scale bar, 100μm. Trend line was plotted between Sox2 and Sox2::HT average nuclear intensity (ANI) distributions of TST18 cells. Correlation coefficient (PMCC, r^2^) is indicated for the trend line.

**Figure 1 – Figure supplement 3.**
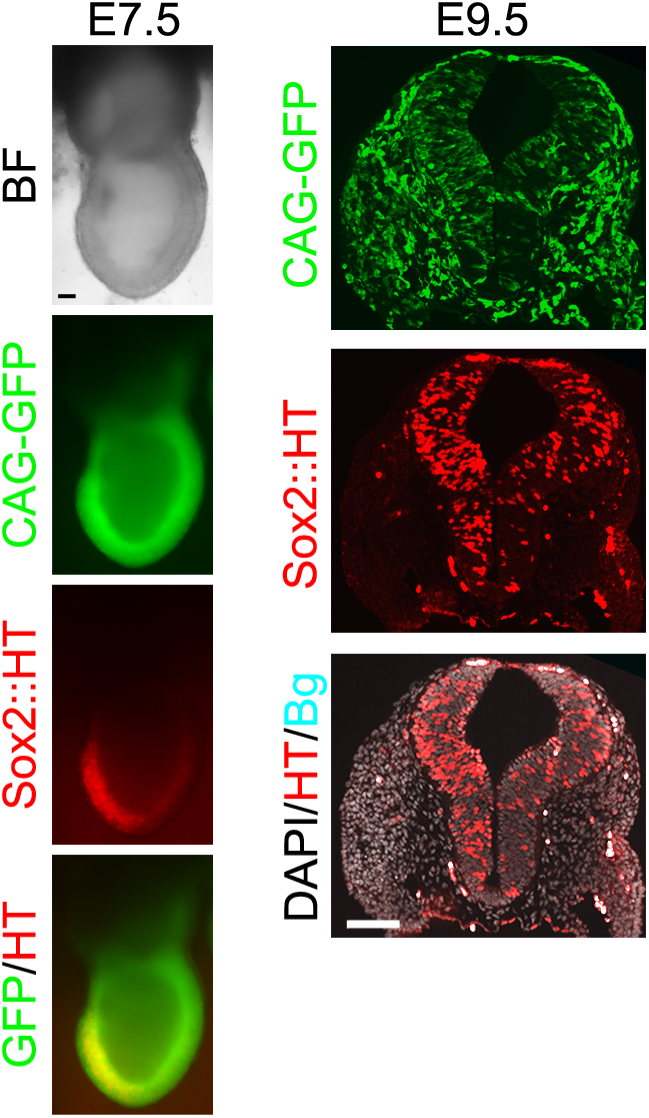
TST (clone 18) cells transfected with a CAG-GFP constitutive reporter were aggregated with isolated morulae and chimeric embryos assessed at the indicated stages for contribution of TST18 cells. Transverse cryostat-sectioning of E9.5 chimeric embryos showed appropriate restriction of Sox2::HT expression. Non-specific auto-fluorescence (cyan) was captured in the far-red (647nm) channel. Scale bar, 100μm.

**Figure 1 – Figure supplement 4.**
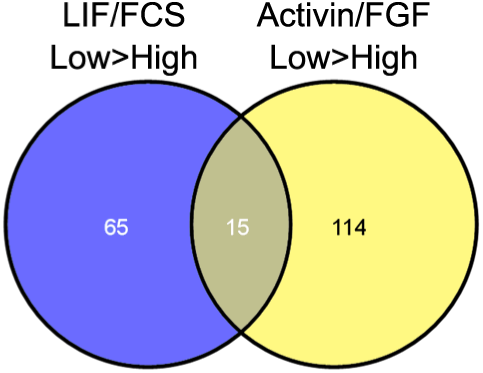
DEGs common to LIF/FCS-low and Activin/FGF-low were not associated to any gene ontology (GO) terms using high-stringency GO term clustering analysis.

**Figure 1 – Figure supplement 5.**
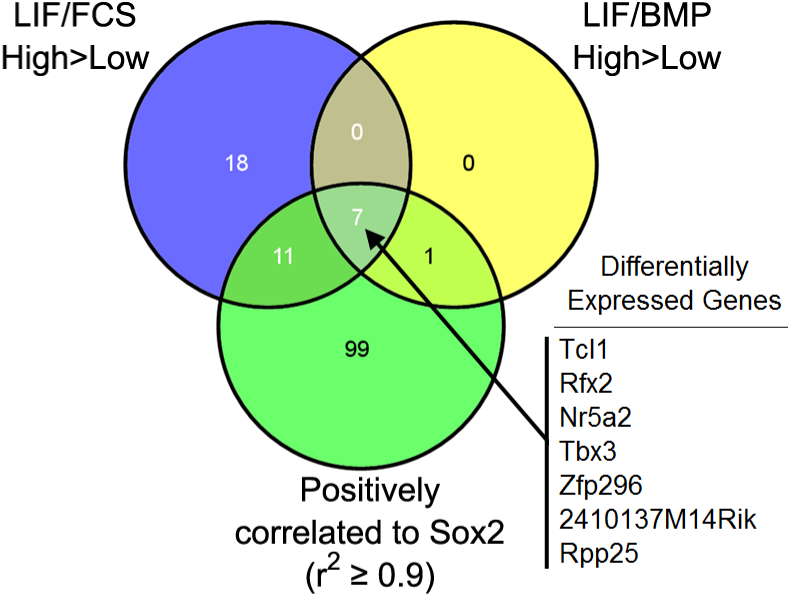
Common differentially expressed genes (DEGs, FDR=0.1) enriched in LIF/FCS-high, LIF/BMP-high and DEGs positively correlated to Sox2 (r^2^ ≥0.9). DEGs listed in descending order of fold-change between LIF/FCS-high and LIF/FCS-low cells.

**Figure 1 – Figure supplement 6.**
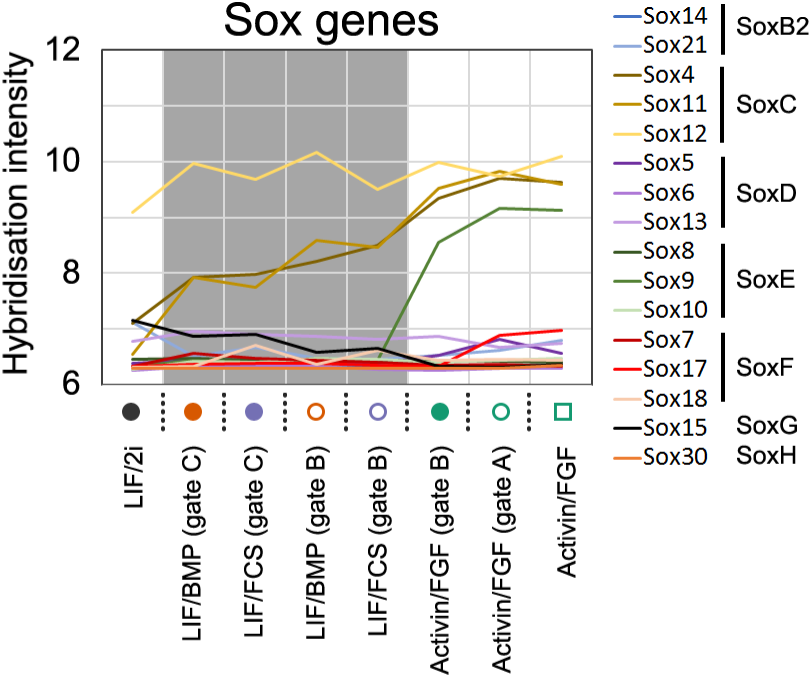
Microarray signal intensity of *Sox* family members in all sorted populations. Sample legend and culture conditions refer to **Figure 1C**.

**Figure 2 – Figure Supplement 1.**
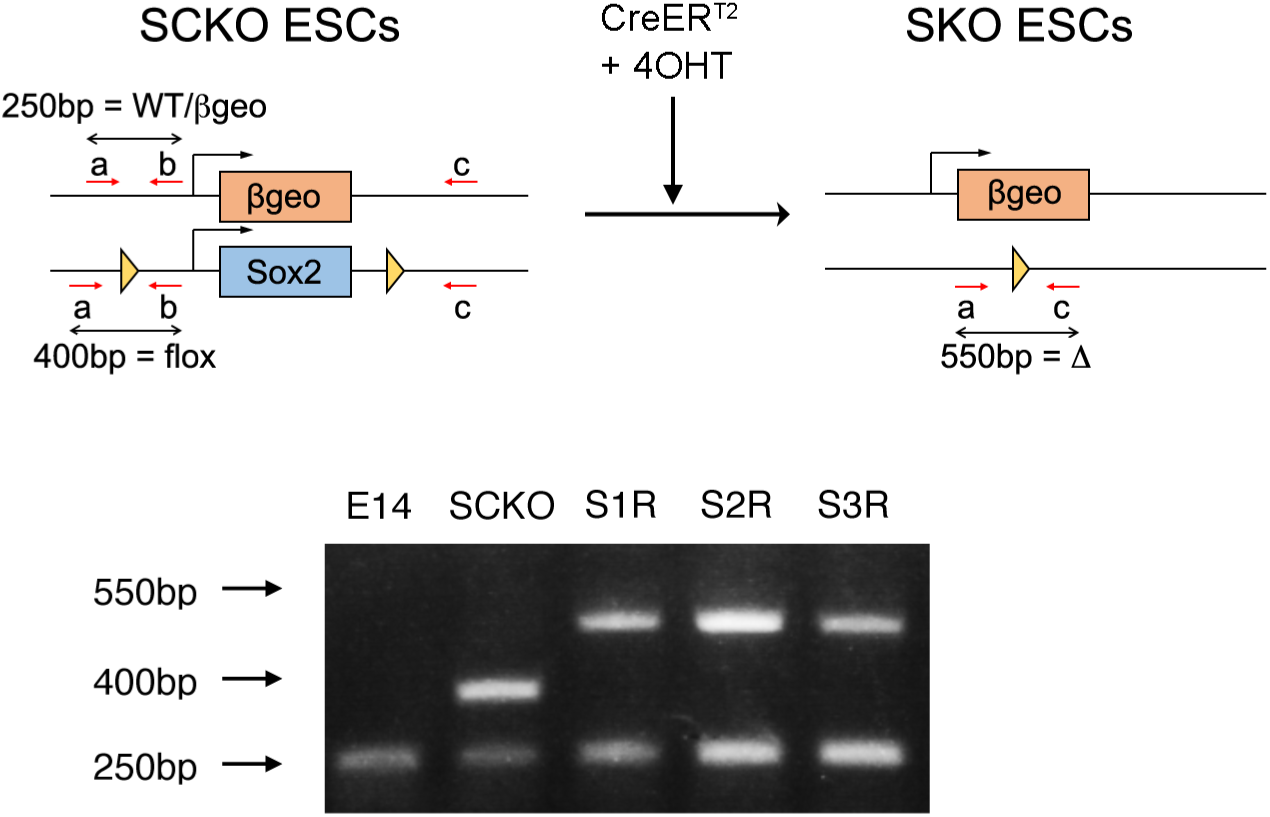
(TOP) Strategy for genotyping the *Sox2* deletion by PCR. Primers a and b flank the position of the *loxP* site 5’ to *Sox2;* primer c flanks the *loxP* site 3’ to *Sox2.* Amplification of wild-type (WT) and βgeo targeted alleles produces a 250bp band whereas the loxP-flanked *Sox2* allele (flox) produces a 400bp band. Upon Cre-mediated recombination, the deleted (Δ) allele produces a 550bp band amplified by primers a+c. (BOTTOM) Wild-type E14Tg2a ESCs (E14) exhibited only the 250bp band. SCKO ESCs exhibited both 250bp and 400bp bands whereas *Sox2*^-/-^ S1R, S2R and S3R ESCs retained the 250bp band and exhibited the 550bp band corresponding to the deleted allele.

**Figure 2 – Figure supplement 2.**
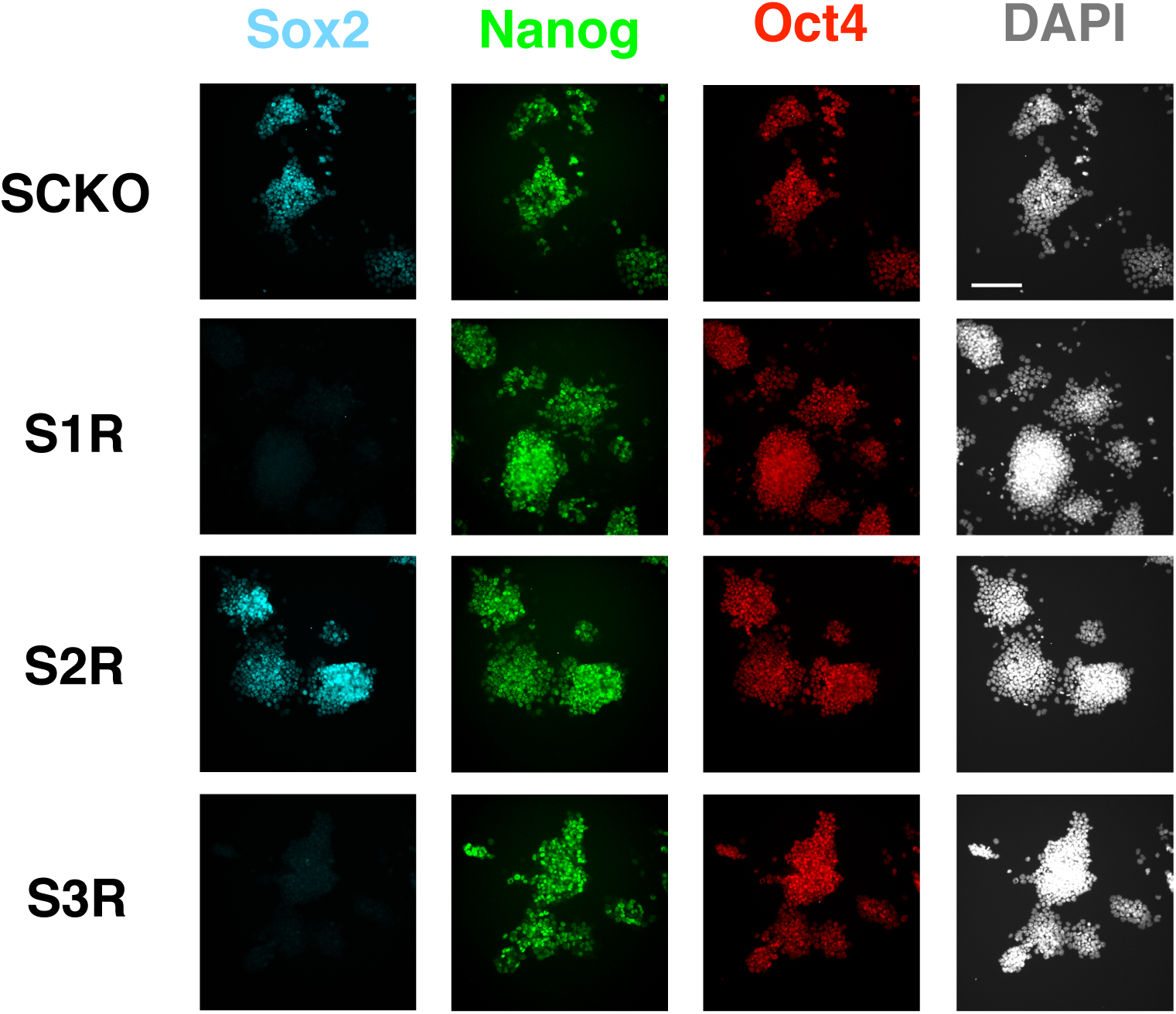
Immunofluorescence analysis of ESCs for Sox2, Nanog and Oct4 in SCKO and S1R, S2R and S3R ESCs. Scale bar, 100μm.

**Figure 3 – Figure supplement 1.**
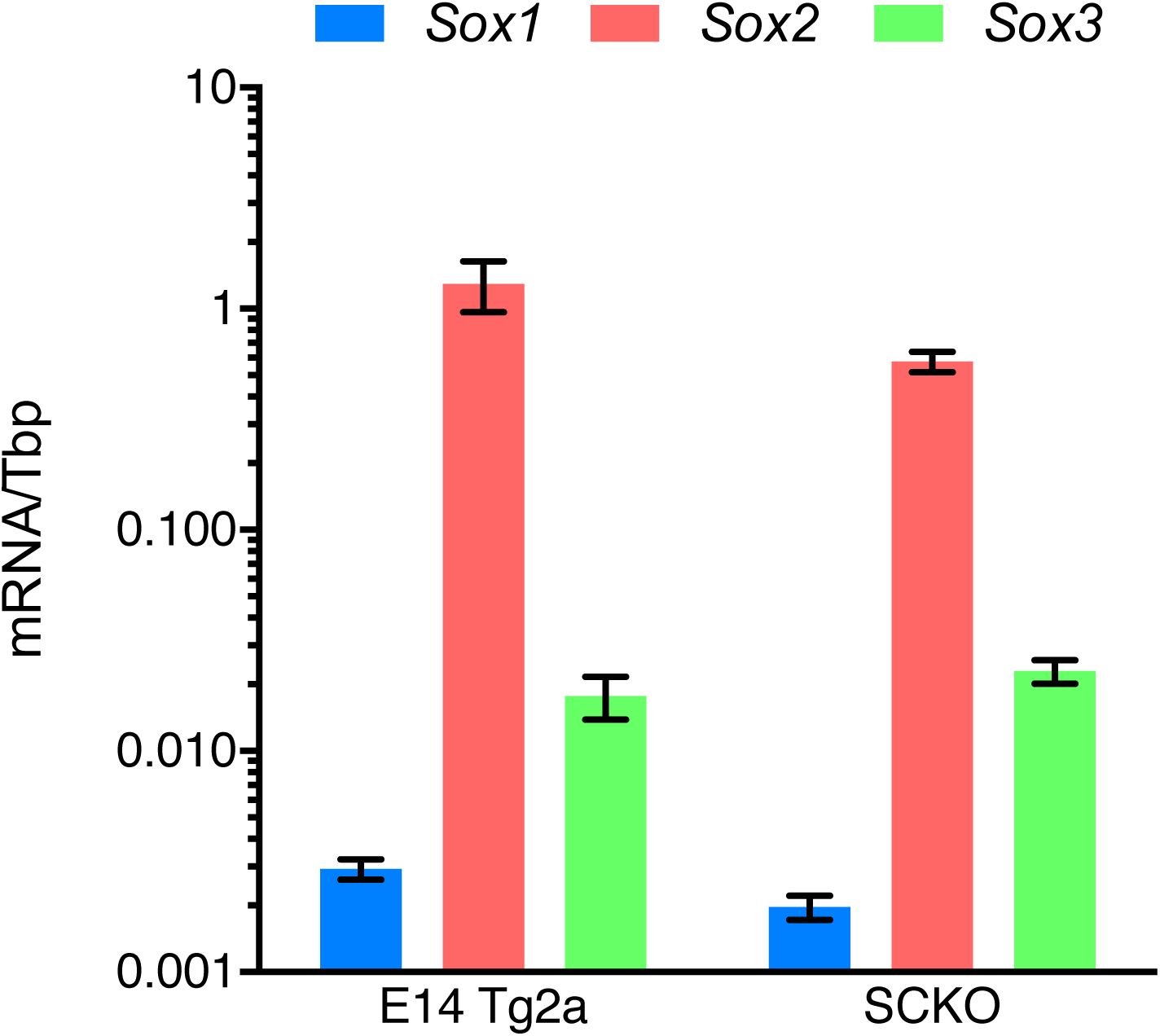
Quantitative transcript analysis showing mRNA levels of the indicated SoxB1 genes in E14Tg2a and SCKO ESCs. mRNA levels were normalised over Tbp. Error bars represent standard error of the mean (n=3).

**Figure 4 – Figure supplement 1.**

Quantitative mRNA analysis showing the levels of the indicated transcripts in E14Tg2a grown in ESC (LIF/FCS) conditions or differentiated in Activin/FGF conditions for two passages (13 days). mRNA levels were normalised over TBP and plotted relatively to ESC (LIF/FCS) values. Error bars represent the standard error of the mean (n=3).

**Figure 5 – Figure supplement 1.**
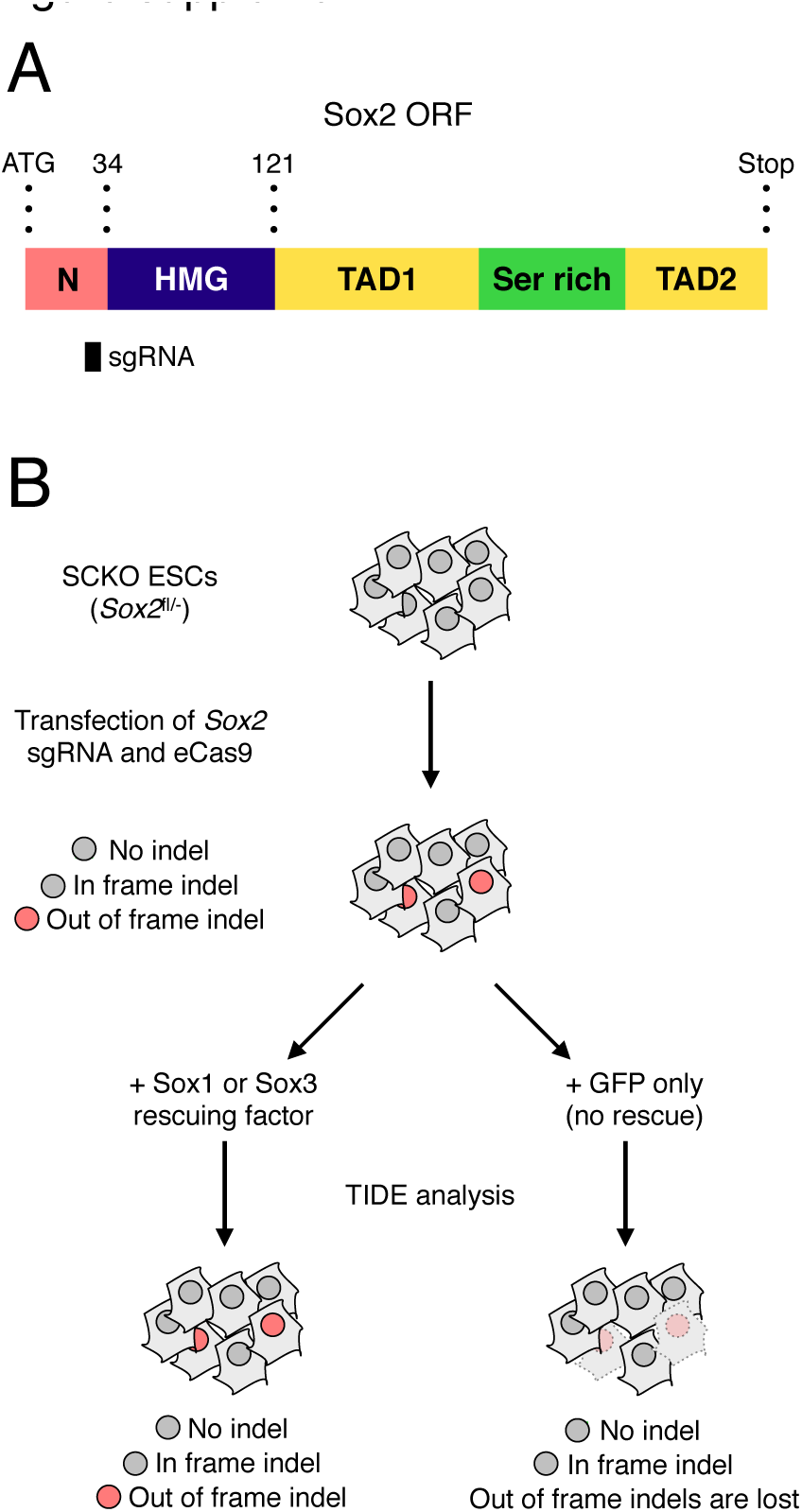
A. Schematic representation of the Sox2 coding sequence (ORF) showing the N-terminal domain (N, pink), the HMG box (dark blue), two transactivation domains (TAD1/2, yellow) and the serine-rich domain (green) (Gagliardi *et al,* 2013). The amino-acid positions of each domain relative to the start (ATG) codon of the ORF are shown. The position of the sgRNA used is represented as black bars. B. Experimental rationale for the expected Sox2 indel pattern in *Sox2^fl/-^* ESCs (SCKO).

**Figure 5 – Figure supplement 2.**
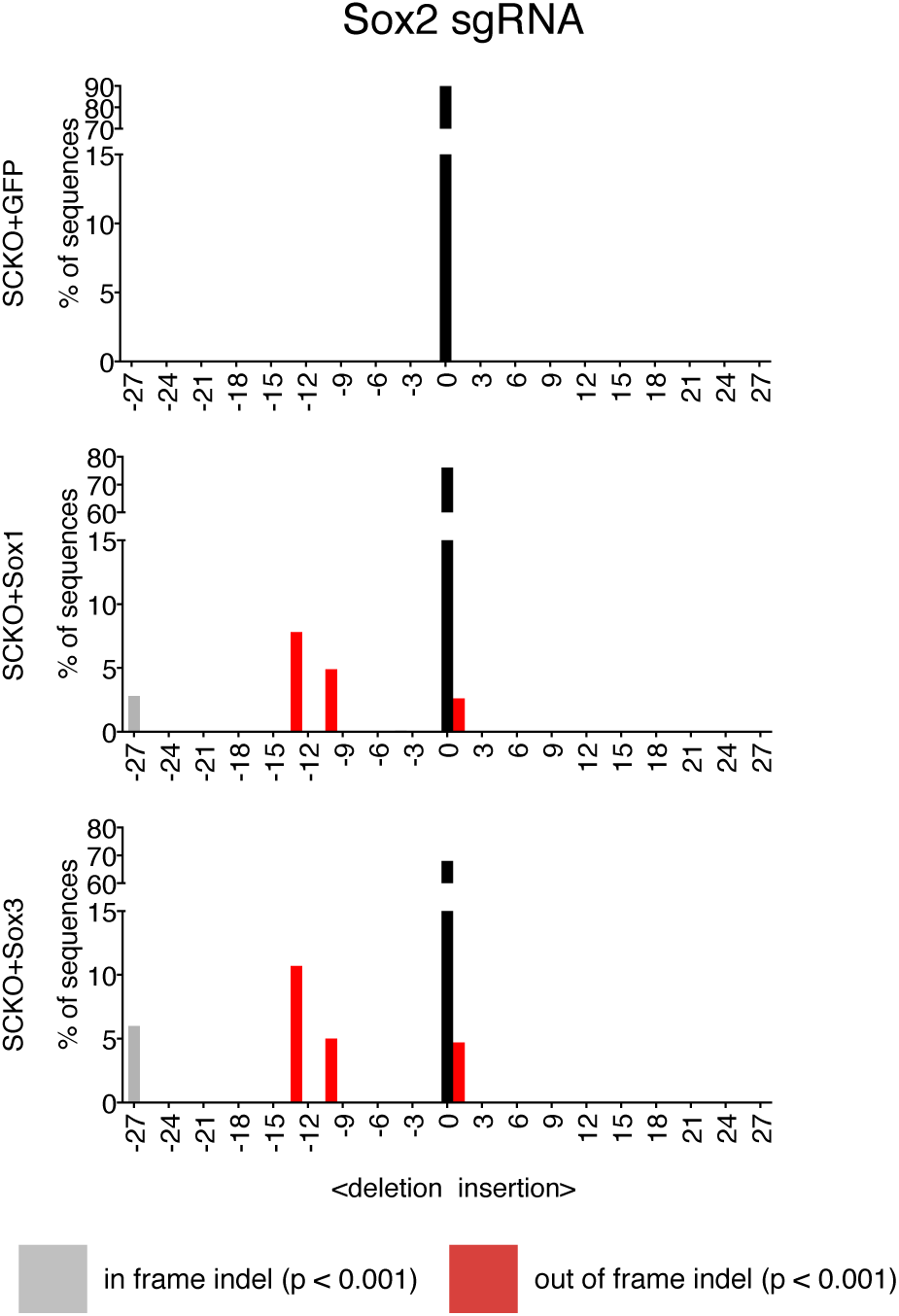
Indel analysis performed using the TIDE tool (https://tide-calculator.nki.nl/) in SCKO. Histograms represent indel frequency and size. Black bars indicate the frequency of unmodified (WT) alleles; grey bars indicate significant in frame indels and red bars indicate significant out of frame indels (p<0.001). Not significant (n/s, p≥0.001) indels are not shown.

**Figure 6 – Figure supplement 1.**
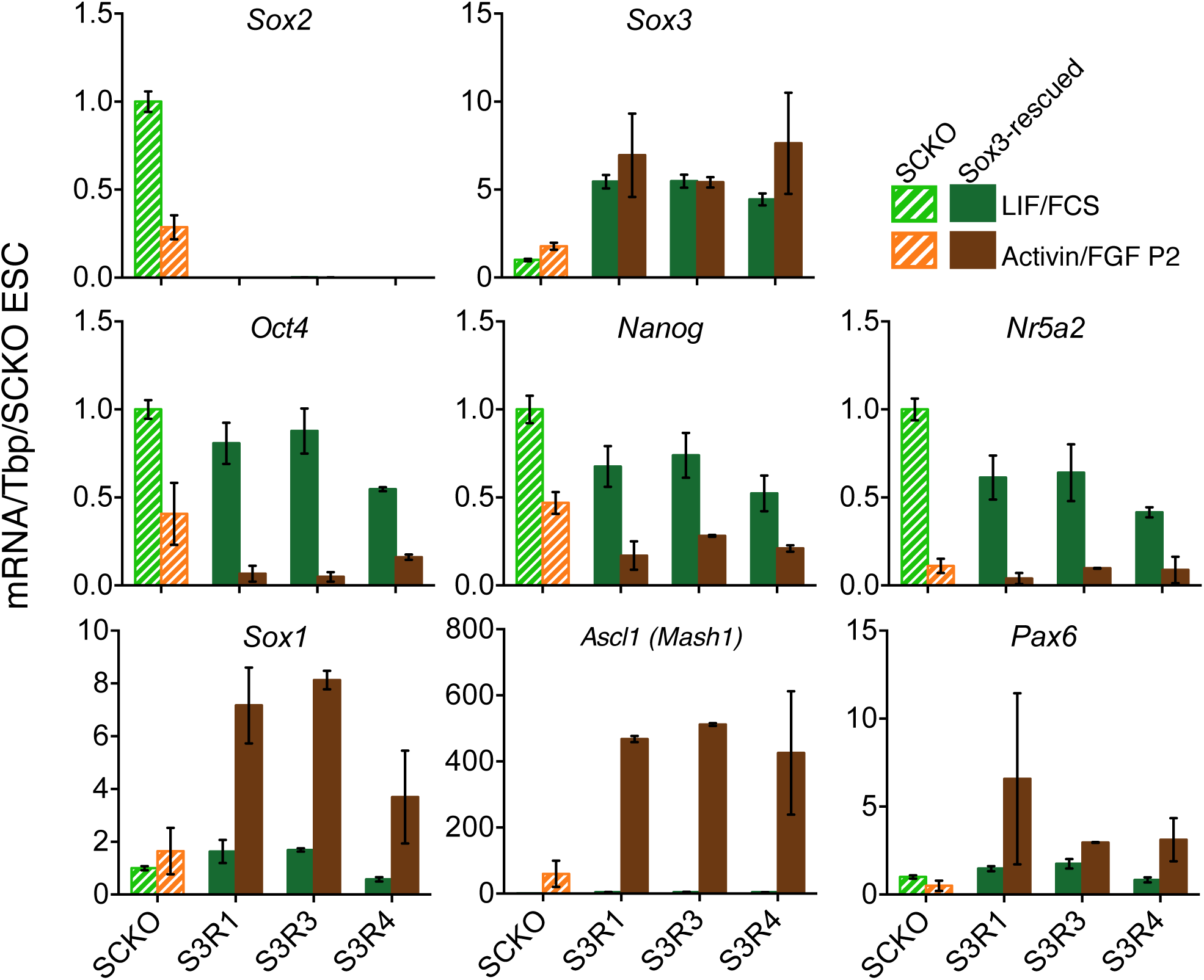
Quantitative mRNA analysis of the mRNA levels of *Sox2, Sox3, Oct4, Nanog* and *Nr5a2* and neural differentiation markers *Sox1, Mash1* and *Pax6* in E14Tg2a cells, *Sox2^fl/-^* cells (SCKO), and *Sox2*^-/-^ S2R and S3R cells maintained in LIF/FCS or differentiated in Activin/FGF conditions (P2). Transcript levels were normalized to TBP and plotted relative to SCKO ESCs. Error bars represent standard error of the mean (n=3 to 5).

**Figure 7 – Figure supplement 1.**
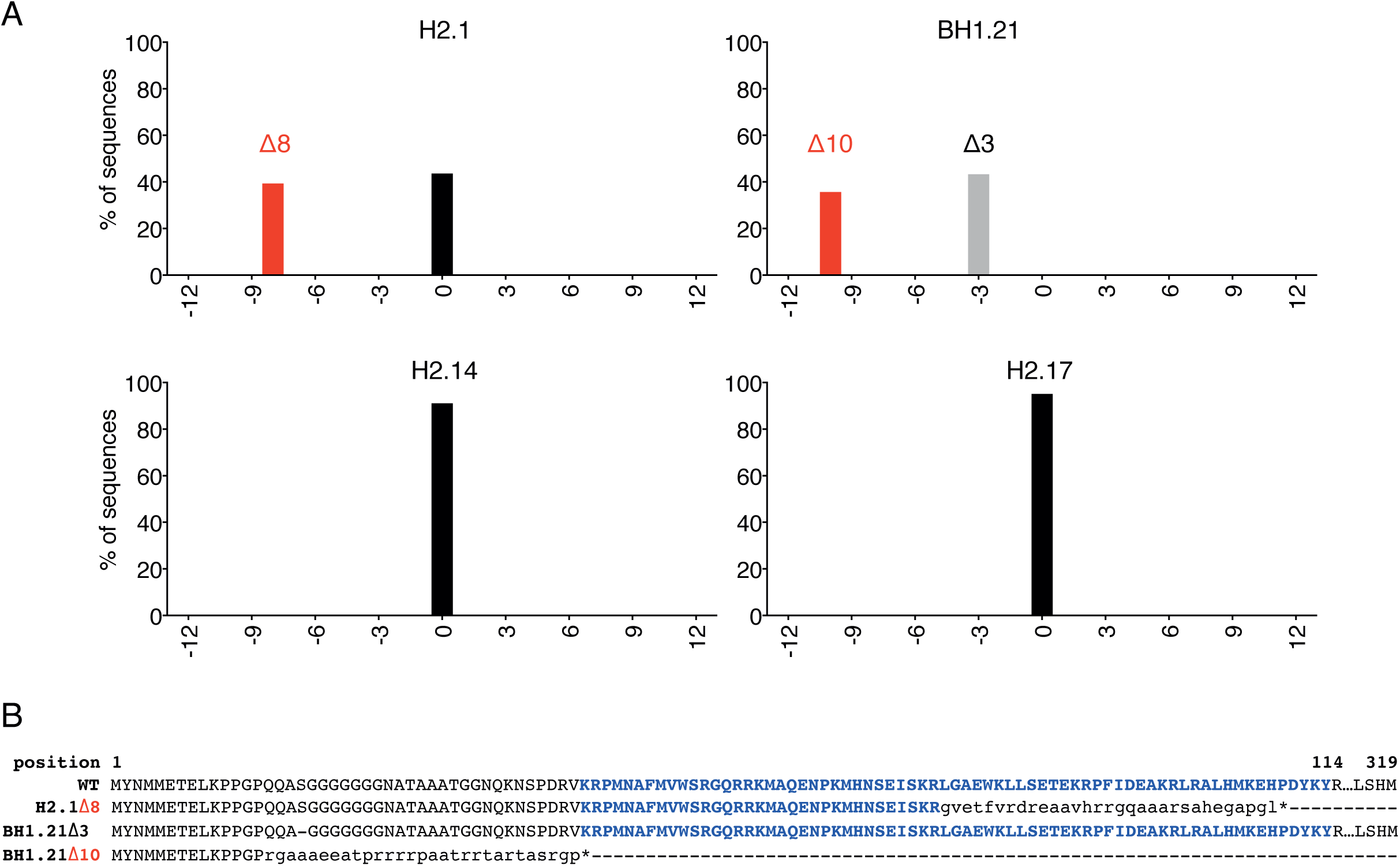
A. Genotyping analysis performed four ESC clones generated after transfection of CRISPR/Cas9 reagents to induce indels within the *Sox2* ORF in E14Tg2a ESCs. PCR amplicons surrounding the Cas9 cutting site were generated starting from gDNA of the individual clones and submitted for Sanger sequencing. Electropherograms containing possible indels were then analysed using the TIDE tool (https://tide-calculator.nki.nl/). Histograms represent frequency and size of deletions found within the *Sox2* alleles of the indicated clones. Black bars indicate the frequency of unmodified (WT) alleles; grey bars indicate in frame indels and red bars indicate out of frame indels. Clone H2.1 carries an out of frame 8-bp (Δ8) on one allele and no deletion on the other allele; clone BH1.21 carries an out of frame 10-bp (Δ10) on one allele and an in frame 3-bp deletion (Δ3) on the other allele; two clones that did not carry any deletion on either *Sox2* allele (H2.14 and H2.17) were also isolated as controls. B. WT SOX2 aminoacid sequence aligned with the sequences produced by the mutated *Sox2* alleles in clones H2.1 and BH1.21 (as indicated). Bold blue font indicates residues forming the HMG DNA binding domain of the protein; small font indicates mutated residues and * indicates premature stop codons generated by the alleles containing out of frame indels; – indicates the deleted serine (S) residue produced by the BH1.21 Δ3 allele.

**Figure 8 – Figure supplement 1.**
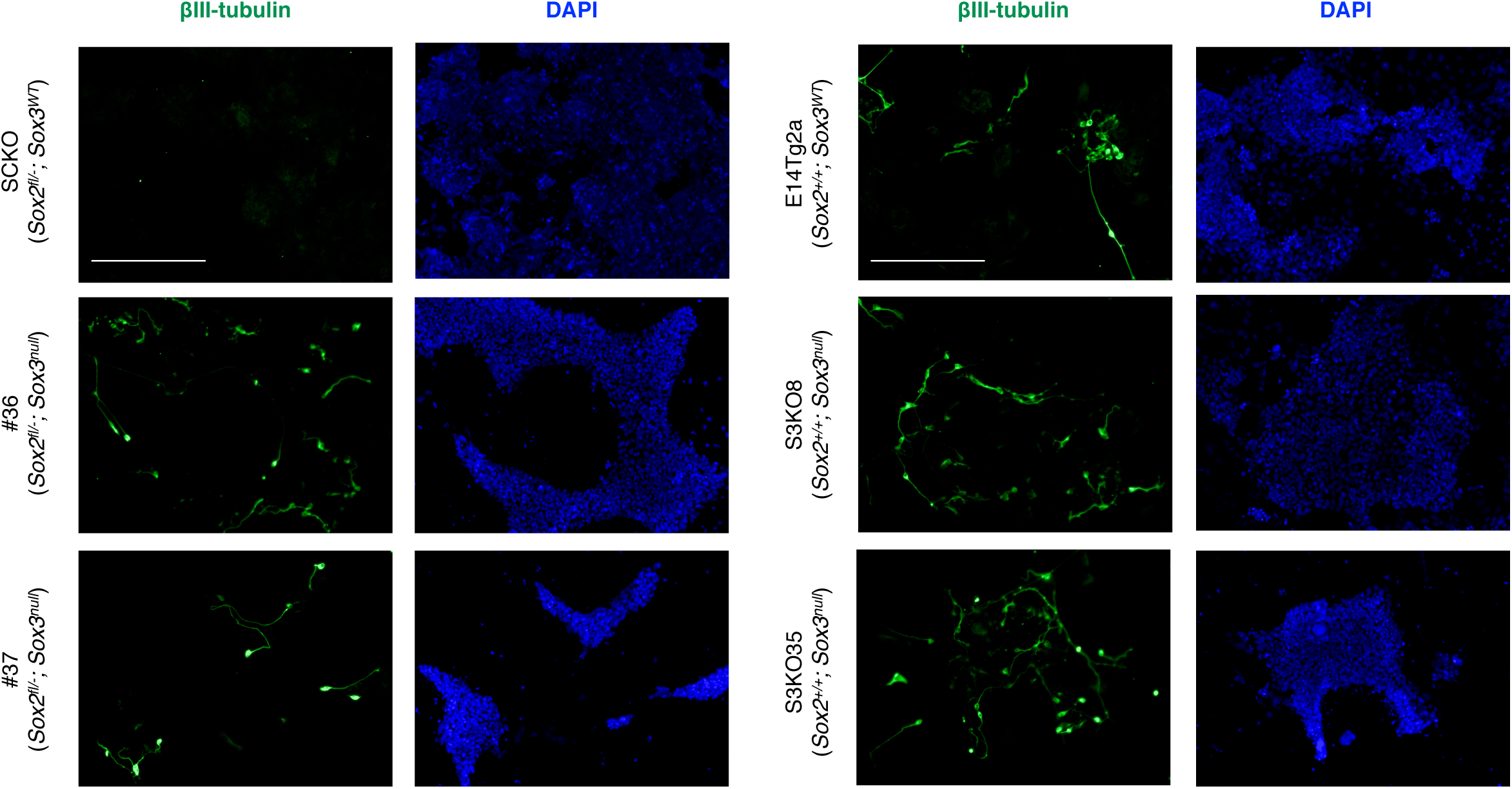
Immunofluorescence staining of the neural marker βIII-tubulin (TUJ1, green) in the indicated cells grown for 96 hours in neural conditions (N2B27 medium); DAPI staining represented in blue. Scale bar, 100μm.

**Figure 8 – Figure supplement 2.**
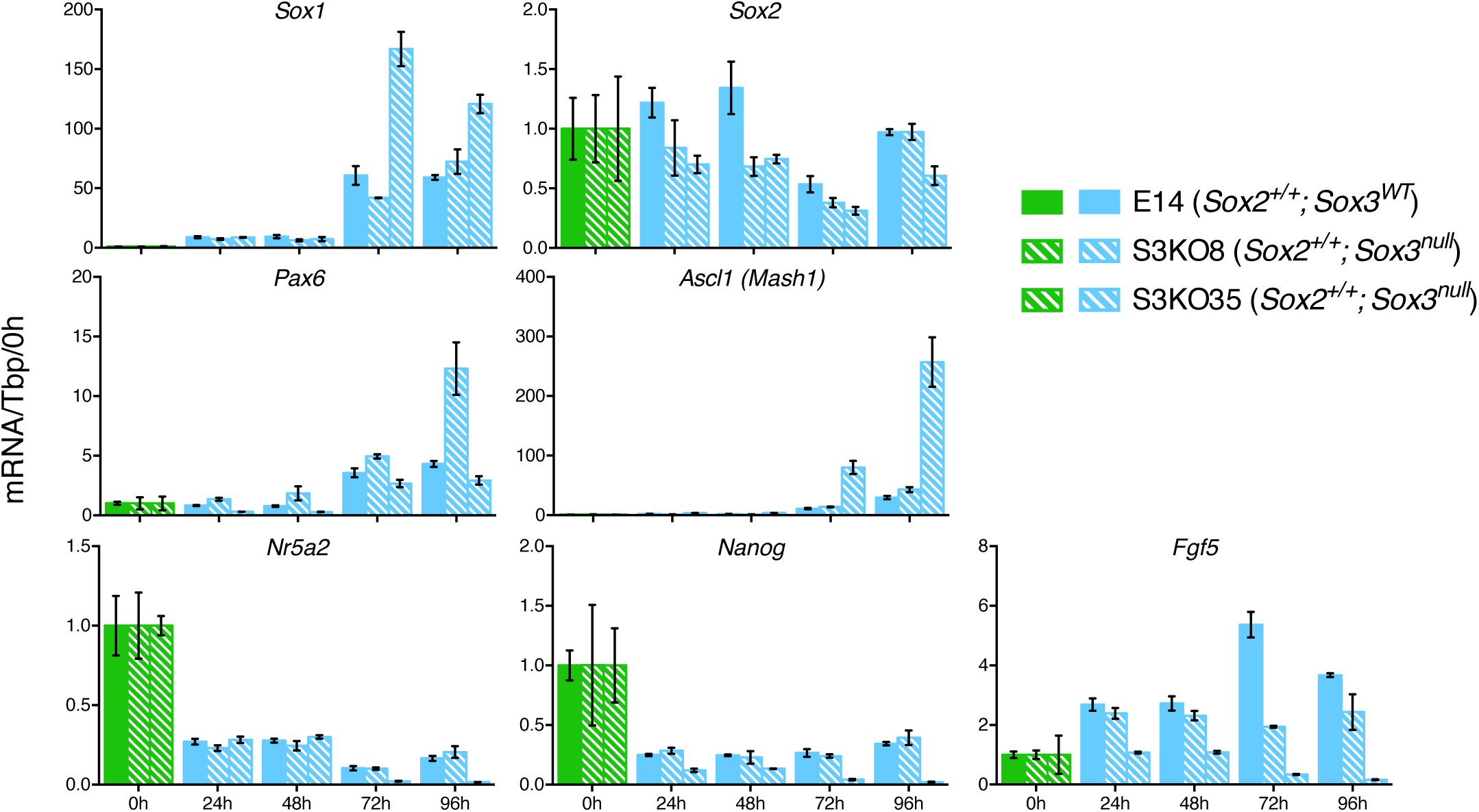
Quantitative mRNA analysis of the indicated transcripts in wild type E14Tg2a cells (*Sox2*^+/+^; *Sox3^WT^*) and in two *Sox2*^+/+^; *Sox3^null^* clones (S3KO8 and S3KO35) (see Figure 3) grown in LIF/FCS conditions and neural differentiation medium (N2B27) for the indicated number of hours. mRNA levels were normalised to TBP and plotted relative to ESC (LIF/FCS). Error bars indicate the standard error of the mean (n=3).

**Supplementary File 1**

Microarray gene expression data described in **Figure 1.**

**Supplementary File 2**

List of the cell lines used in this study with their genotype and additional transgenes if present.

**Supplementary File 3**

List of the PCR primers and sgRNAs used in this study.

